# Divergent role of CD8 T cells with distinct metabolic phenotypes during curative radio-immunotherapy in hot versus cold tumors

**DOI:** 10.1101/2025.10.24.684274

**Authors:** Alexa R. Heaton, Nathaniel J. Burkard, Anqi Gao, Anna Hoefges, Arika S. Feils, Samantha K. Burkard, Mildred A. Felder, Noah W. Tsarovsky, Daniel V. Spiegelman, Garrett M. Lublin, Alina A. Hampton, Aurora D’Amato, Huy Q. Dinh, Alexander L. Rakhmilevich, Amy K. Erbe, Paul M. Sondel, Melissa C. Skala

**Affiliations:** Morgridge Institute for Research, Madison, WI, USA; Department of Human Oncology, University of Wisconsin, Madison, WI, USA; Department of Biomedical Engineering, University of Wisconsin, Madison, WI, US; McArdle Laboratory for Cancer Research, University of Wisconsin, Madison, WI, USA; Department of Neuroscience, University of Wisconsin, Madison, WI, USA; Department of Computer Sciences, University of Wisconsin, Madison, WI, USA; Global Health Institute, University of Wisconsin, Madison, WI, USA; Department of Biostatistics and Medical Informatics, University of Wisconsin, Madison, WI, USA; Department of Pediatrics and Department of Human Oncology, University of Wisconsin, Madison, WI, USA

## Abstract

Immunotherapy has potential for impactful cancer cures by empowering patients’ own immune cells. We developed a radio-immunotherapy regimen that can cure large immunologically hot and cold murine tumors. Here, we explored the divergent role of CD8 T cells during this radio-immunotherapy in contrasting hot colon carcinoma versus cold melanoma. We introduced an immunocompetent mouse model with mCherry-expressing CD8 T cells to provide cell tracking *in vivo*. We investigated single-cell function, metabolism, and gene expression temporal changes using flow cytometry, *in vivo* multiphoton imaging, single-cell RNA sequencing, and multiplexed immunofluorescence to determine the underlying mechanisms. We found that in contrast to the hot colon carcinoma model, CD8 T cells from the cold melanoma model do not drive tumor cures, despite getting activated, possibly due to a static oxidative metabolism and exhausted phenotype plus down regulation of tumor MHC-I expression. These findings have implications for improving immunotherapy response in immunologically cold cancers.

## INTRODUCTION

Despite recent immunotherapy progress, the majority of cancer patients are not yet benefiting.^1–3^ To improve response, we and others have focused on combining immunotherapies with synergistic immunomodulatory radiation therapy that produces durable responses across many solid tumors.^3–12^ This mirrors clinical experience where combination therapy including immune checkpoint inhibitors (ICI) has improved survival.^3,13–15^ Our recent preclinical work has treated an immunologically cold murine melanoma where immune infiltration is limited, major histocompatibility complex class I (MHC-I) expression is very low, and there is no response to ICI – though our radio-immunotherapy regimen is effective.^9,11,16–19^ Here, we characterize the immune response and determine underlying mechanisms that drive this curative radio-immunotherapy combination.

Within immuno-oncology, over 80% of immunotherapy products in clinical trials work via CD8 T cells.^1,20–23^ In the tumor microenvironment (TME), numerous barriers can prevent anti-tumor CD8 T cell function, including physical barriers (dense extracellular matrix and abnormal vasculature that exclude tumor infiltration^24–27^), immuno-suppressive cells (tumor cells, fibroblasts, myeloid derived suppressor cells, and others that dampen immune function ^27–40^), and metabolic barriers (competition for oxygen and nutrients in theTME^41–50^). Metabolic shifts often precede functional shifts and have been correlated to immune cell role and phenotype – providing a window into CD8 T cell function.^45,50–58^ There is a need for methods that can quantify these metabolic shifts in living animals as this may help develop more effective immunotherapy by providing a deeper understanding of tumor and immune cell metabolism during tumorigenesis and treatment. Optical metabolic imaging (OMI) of endogenous metabolic co-enzymes can address this issue by providing label-free metabolic readouts in living animals.^59,60^

We have established label-free OMI to probe single-cell function, phenotype, and therapeutic response within tumors in living animals. OMI relies on autofluorescence from NAD(P)H and FAD, which are electron shuttles that drive most metabolic processes in the cell.^61,62^ We can image and quantify intensity-based changes – related to the concentration of each molecule within the cell – and lifetime-based changes – defined as the time it takes for a molecule to relax back to the ground state after photon excitation, which informs on protein binding activity. Protein-bound FAD and free NAD(P)H have a shorter lifetime (τ_1_) while free FAD and protein-bound NAD(P)H have a longer lifetime (τ_2_). The fractional contribution of coenzyme in each conformation is denoted by α_1_ and α_2_.^63–66^ This technology has provided insights into cancer and immune cell metabolism including metabolic differences between healthy and cancerous tissues,^59,67^ and immune cell metabolic shifts during activation^60,68–70^ and immunotherapy.^71^ We now expand on this work by performing *in vivo* tumor metabolic imaging during curative radio-immunotherapy.

Here, we investigated the therapeutic mechanisms of a curative radio-immunotherapy combination on two contrasting tumor models: immunologically hot MC38 colon carcinoma and immunologically cold B78 melanoma. We determined which immune cell populations drive tumor response and temporal changes within the TME during treatment, with a focus on CD8 T cells. We quantified immune and tumor cell metabolic changes using single-cell label-free OMI in live mouse tumors and correlated these findings with multiparameter flow cytometry and single-cell RNA sequencing. Overall, we employed a multimodal single-cell approach to investigate the divergent roles of CD8 T cells during radio-immunotherapy cures, in hot versus cold tumors, to probe the underlying divergent mechanisms related to immunotherapy response.

## RESULTS

### CD8-mCherry reporter mouse model shows high specificity of mCherry expression to CD8 T cells

To facilitate *in vivo* imaging of dynamic CD8 T cells, we created a CD8-mCherry reporter mouse model where CD8 cells express the red fluorescent protein mCherry (**Extended Data Fig. 1A**). To characterize these mice, CD3+ splenic T cells were isolated from both C57BL/6J wild type and CD8-mCherry reporter mice and the T cells were imaged *in vitro* via OMI. All T cells contained NAD(P)H while only the reporter mouse expressed mCherry in ∼50% of the splenic T cells, the CD8 subset (**Extended Data Fig. 1B**). Additionally, multiplexed immunofluorescence (mIF) images of untreated B78 melanoma tumors and spleens from reporter mice showed that mCherry signal is only expressed on cells that also co-express CD8 (**Extended Data Fig. 1C**), suggesting that the knock-in near the CD8α allele was specific with no off-target expression. Flow cytometry of B78 tumors and spleens from untreated reporter mice showed the majority of CD8 cells (∼94%) were positive for mCherry expression while other immune cells expressed minimal mCherry (∼1%) (**Extended Data Fig. 1D**).

### MC38 colon carcinoma is highly responsive to radio-immunotherapy regimen

MC38 is a well-studied, immunologically hot tumor with high mutational burden.^72,73^ CD8-mCherry reporter mice bearing ∼150 mm^3^ intradermal^74^ (I.D.) flank MC38 tumors were treated with external beam radiation therapy, αCTLA-4, and IL-2 (**Fig. 1A**). Our radio-immunotherapy combination induced complete responses in 91% of treated mice (**Fig. 1B-C**) and significantly increased survival (**Fig. 1D**). When cured mice were rechallenged with a second MC38 tumor they cleared 100% of the rechallenged tumors while naïve mice cleared 0%, indicating this radio-immunotherapy produces a strong immune memory response (**Fig. 1E**). This radio-immunotherapy efficacy was also reproduced in C57BL/6J mice to ensure the response was not specific to our reporter mice (**Extended Data Fig. 2A-D**).

**Figure 1.**
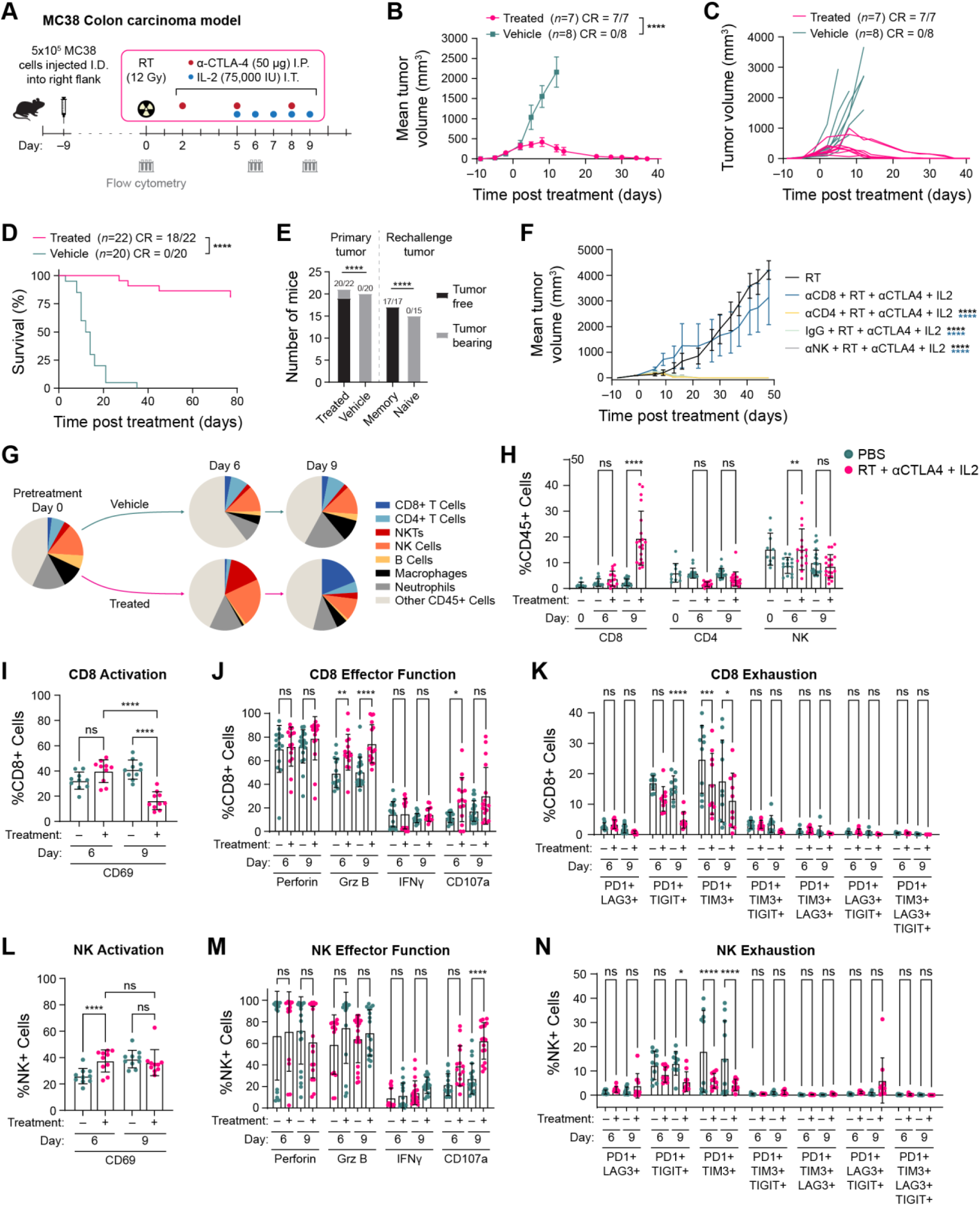
Radio-immunotherapy leads to strong responses and immune infiltrate changes in MC38 colon carcinoma. **A)** Experimental timeline with flow cytometry. **B)** Mean tumor-growth curves for treated and vehicle MC38-bearing mice (pairwise contrasts, *n*=7-8 mice/group, mean ± SEM, 1 representative experiment shown, 3 experimental repeats). **C)** Individual tumor-growth curves for treated and vehicle MC38-bearing mice (*n*=7-8 mice/group, 1 representative experiment shown, 3 experimental repeats). **D)** Kaplan-Meier survival curve of treated and vehicle MC38-bearing mice (pairwise log-rank test, *n*=20-22 mice/group, 3 experimental repeats combined). **E)** Number of tumor free mice from treated and vehicle primary tumors and from treated and naïve tumor rechallenge (Fisher exact test, primary *n*=20-22, rechallenge *n*=15-17 mice/group, 3 experimental repeats). **F)** Mean tumor-growth curves during CD8, CD4, and NK cell depletion of MC38-bearing mice (*n*=5 mice/group, pairwise contrasts, 1 representative experiment shown, 2 experimental repeats, mean ± SEM). Note green, yellow, and gray curves are superimposed on the X-axis. **G)** Flow cytometry of immune cell subsets in tumors from treated and vehicle MC38-bearing mice across treatment time (*n*=5-10 mice/timepoint, 2 experimental repeats, %CD45+ cells). **H)** Flow cytometry of CD8, CD4, NK cell changes from treated and vehicle MC38-bearing mice across treatment time (one-way ANOVA, *n*=5-10 mice/group, 2-3 experimental repeats, mean ± SD, %CD45+ cells). **I-K)** Flow cytometry of CD8 and **L-N)** NK activation, effector function, and exhaustion from treated and vehicle MC38-bearing mice across treatment time (*n*=10-20 mice/group, one-way ANOVA, mean ± SD, %CD8+ or %NK+ cells, 2-4 experimental repeats) (\**p* < 0.05, \*\**p* < 0.01, \*\*\**p* < 0.001, \*\*\*\**p* < 0.0001).

### Hot MC38 colon carcinoma cures are driven by CD8 T cells

To probe which immune cells are driving these MC38 responses, we selectively depleted CD8, CD4, or NK cells. Radio-immunotherapy treated MC38-bearing mice depleted of CD4 or NK cells behaved no differently than treated mice receiving control IgG antibody – complete responses were observed in all mice (**Fig. 1F**). In contrast, radio-immunotherapy treated mice depleted of CD8 cells lost the ability to clear their tumors and responded similarly to mice that received radiation therapy alone but significantly different than αCD4, αNK, IgG groups (**Fig. 1F**). Thus, CD8 cells are critical for MC38 tumor response to our radio-immunotherapy *in vivo*. We further explored immune function during radio-immunotherapy using multi-parameter flow cytometry on day 0, 6, and 9 (gating strategies in **Extended Data Fig. 3A)**. We showed that tumor infiltrating immune populations changed over time and with therapy, including a significant expansion of NK cells on day 6, a significant reduction of neutrophils on day 9, and a significant expansion of CD8 cells on day 9 (**Fig. 1G-H**, **S3B**). Functional measurements showed that CD8 cells from treated mice are more activated on day 6 versus day 9 (CD69 expression) (**Fig. 1I**), contain significantly higher granzyme B on both days, express significantly higher levels of degranulation (CD107a) on day 6 (**Fig. 1J**), and express significantly less exhaustion markers compared to vehicle mice (**Fig. 1K**). As we saw an expansion of NK cells during therapy, we also probed their functional changes and found that NK cells from treated mice were significantly more activated on day 6 (**Fig. 1L**) but showed minimal differences in effector function except for significantly increased degranulation (CD107a) on day 9 when compared to vehicle mice (**Fig. 1M**) and expressed significantly less exhaustion markers compared to vehicle mice (**Fig. 1N**). CD4 cells from treated mice were significantly more activated on day 6 compared to day 9 treated mice, contained significantly higher cytolytic granules (perforin and granzyme B), expressed higher CD107a, and expressed significantly less exhaustion markers compared to vehicle mice (**Extended Data Fig. 3C-E**). In summary, antibody depletion experiments and a multi-parameter flow cytometry time-course show that CD8 T cells drive MC38 tumor cures though some effector function is also observed within NK and CD4 T cell populations in this hot tumor.

### In vivo imaging captures MC38 CD8 T cell metabolic dynamics from early glycolytic state to later memory state

We performed *in vivo* label-free OMI to track single CD8 T cell metabolic changes across treatment on days 0, 6, and 9 (**Fig. 2A**). Representative fluorescence intensity images showed increased CD8 infiltration and cell size (white outlines mark CD8 T cells) in treated mice versus vehicle mice (**Fig. 2B**, top row). Representative NAD(P)H α_1_ and FAD α_1_ images showed that CD8 T cell NAD(P)H and FAD protein-binding changes over treatment time-course (**Fig. 2B**, middle and bottom rows). Quantified single CD8 T cell metabolic changes showed an initial trend towards increased NAD(P)H α_1_ on day 6, suggesting a glycolytic shift in metabolism, followed by a significant decrease on day 9, suggesting an oxidative shift in metabolism (**Fig. 2C**). Furthermore, CD8 T cell metabolic changes showed a significant decrease in FAD α_1_ on day 6, also suggesting a glycolytic shift in metabolism, followed by an increase on day 9, also suggesting an oxidative shift in metabolism (**Fig. 2D**). Additionally, OMI showed that on day 6 the CD8 T cell size increased, often correlated to T cell activation and proliferation, but not on day 9 (**Fig. 2E**). We trained a binary classifier on CD8 T cell *in vivo* OMI variables and tested predictive accuracy for each CD8 T cell originating from a treated versus vehicle mouse tumor. On day 6, the classifier identified CD8 T cells from treated versus vehicle mice with high accuracy (area under the curve (AUC) = 0.94, **Fig. 2F**).

**Figure 2.**
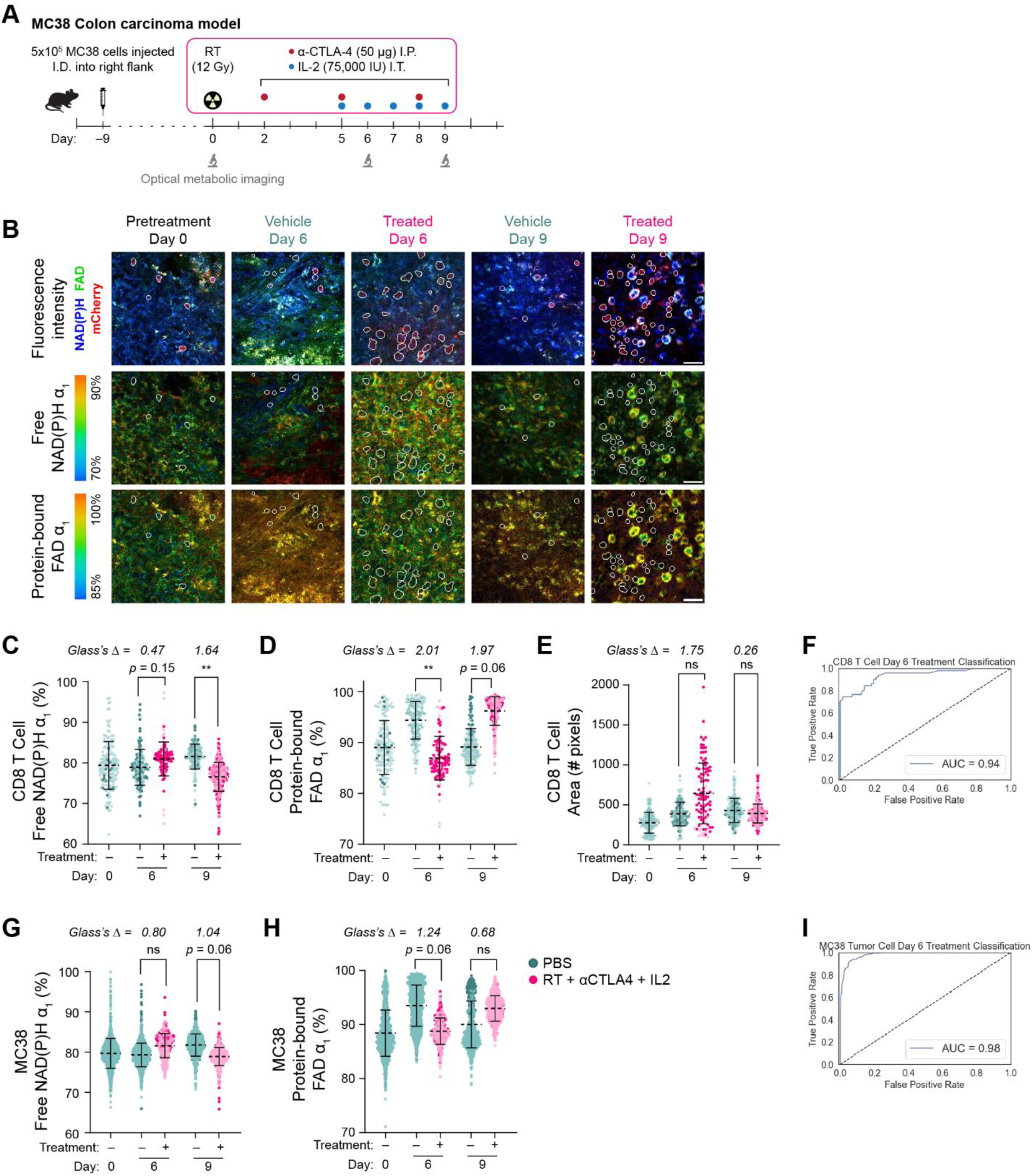
In vivo fluorescence intensity and lifetime imaging of CD8 T cells in MC38 colon carcinoma tumors shows metabolism changes with treatment. **A)** Experimental timeline with *in vivo* OMI. **B)** Representative *in vivo* MC38 colon carcinoma tumor images across time. Fluorescence intensity images (top row) show infiltrating CD8 T cells (red) expressing mCherry and autofluorescent NAD(P)H (blue) and FAD (green) present in all cells. NAD(P)H α_1_ (middle row) and FAD α_1_ (bottom row) images show spatial distribution of metabolic changes within the tumors. CD8 T cell outlines (white) overlaid on all images for clarity (1 experiment shown, 2 experimental repeats, scale bar 50 μm). **C-E)** CD8 T cell NAD(P)H α_1_ (proportion of free NAD(P)H), FAD α_1_ (proportion of protein-bound FAD), and area changes during therapy. **F)** Receiver operating characteristic (ROC) curve (area under the curve (AUC) = 0.94) classifying treated or vehicle CD8 T cells from day 6. **G-H)** MC38 tumor cell NAD(P)H α_1_ and FAD α_1_ changes during therapy. **I)** ROC curve (AUC = 0.98) classifying treated or vehicle MC38 tumors cells from day 6. **C-E)** *n*=4 mice/day, *n*=2-4 mice/group, Day 0 *n*=152 cells, Day 6 *n*=244 cells, Day 9 *n*=469 cells, CD8 T cells imaged, mixed-effects ANOVA and Glass’s Δ, mean ± SD, 2 experimental repeats. **G-H)** *n*=4 mice/day, *n*=2-4 mice/group, Day 0 *n*=1445 cells, Day 6 *n*=1882 cells, Day 9 *n*=1409 cells, MC38 tumor cells imaged, mixed-effects ANOVA and Glass’s Δ, mean ± SD, 2 experimental repeats. **F, I)** Random Forest algorithm, 50% train:50% test; **Extended Data Table 1** shows 9 variables included, day 0 data not included.

### In vivo imaging captures MC38 tumor cell metabolic dynamics during radio-immunotherapy

We also captured single MC38 tumor cell metabolic changes across treatment (**Fig. 2A**). Representative fluorescence intensity images showed changes in TME over time, with lower tumor density later in treatment (**Fig. 2B**, top row). Representative NAD(P)H α_1_ and FAD α_1_ images showed that MC38 tumor cell NAD(P)H and FAD protein-binding changes over treatment time-course (**Fig. 2B**, middle and bottom rows). Single cell MC38 metabolic changes showed an initial, insignificant increase in NAD(P)H α_1_, suggesting a glycolytic shift in metabolism, on day 6 followed by a decrease on day 9, indicating an oxidative shift in metabolism (**Fig. 2G**). In contrast, MC38 tumor cell metabolic changes showed a decrease FAD α_1_ on day 6, also suggesting a glycolytic shift in metabolism, with no change on day 9 (**Fig. 2H**). These metabolic shifts were validated by plating MC38 *in vitro* and applying known metabolic inhibitors including an electron transport chain complex IV inhibitor, sodium cyanide (NaCN), and a glycolysis inhibitor, 2-Deoxy-D-glucose (2DG) (**Extended Data Fig. 4B-E**). We trained a binary classifier on *in vivo* MC38 tumor cell metabolic variables and tested sensitivity to treatment vs. vehicle condition. On day 6, the classifier identified MC38 tumor cells from treated versus vehicle mice with high accuracy (AUC = 0.98, **Fig. 2I**).

### B78 melanoma responsive to radio-immunotherapy regimen

Next, we transitioned to a difficult to treat, immunologically cold B78 melanoma tumor. CD8-mCherry reporter mice bearing I.D. ∼150mm^3^ flank B78 tumors were treated with radiation therapy, αCTLA-4, and Hu14.18-IL2 immunocytokine (**Fig. 3A**). Our radio-immunotherapy combination induced complete responses in 58% of treated mice (**Fig. 3B-C, E**) and significantly increased survival (**Fig. 3D**). Cryo-fluorescence tomography imaging showed that radio-immunotherapy induced increased numbers of mCherry+ CD8, CD4, and NKT cells within B78 tumors and lymph nodes, compared to control mice (**Extended Data Fig. 5A-C**). When cured mice were rechallenged with a second B78 tumor without additional therapy, they cleared 80% of the rechallenged tumors while naïve mice cleared 0%, indicating this radio-immunotherapy produces a strong immune memory response (**Fig. 3E**). Efficacy was also reproduced in C57BL/6J mice (**Extended Data Fig. 2E-H**).

**Figure 3.**
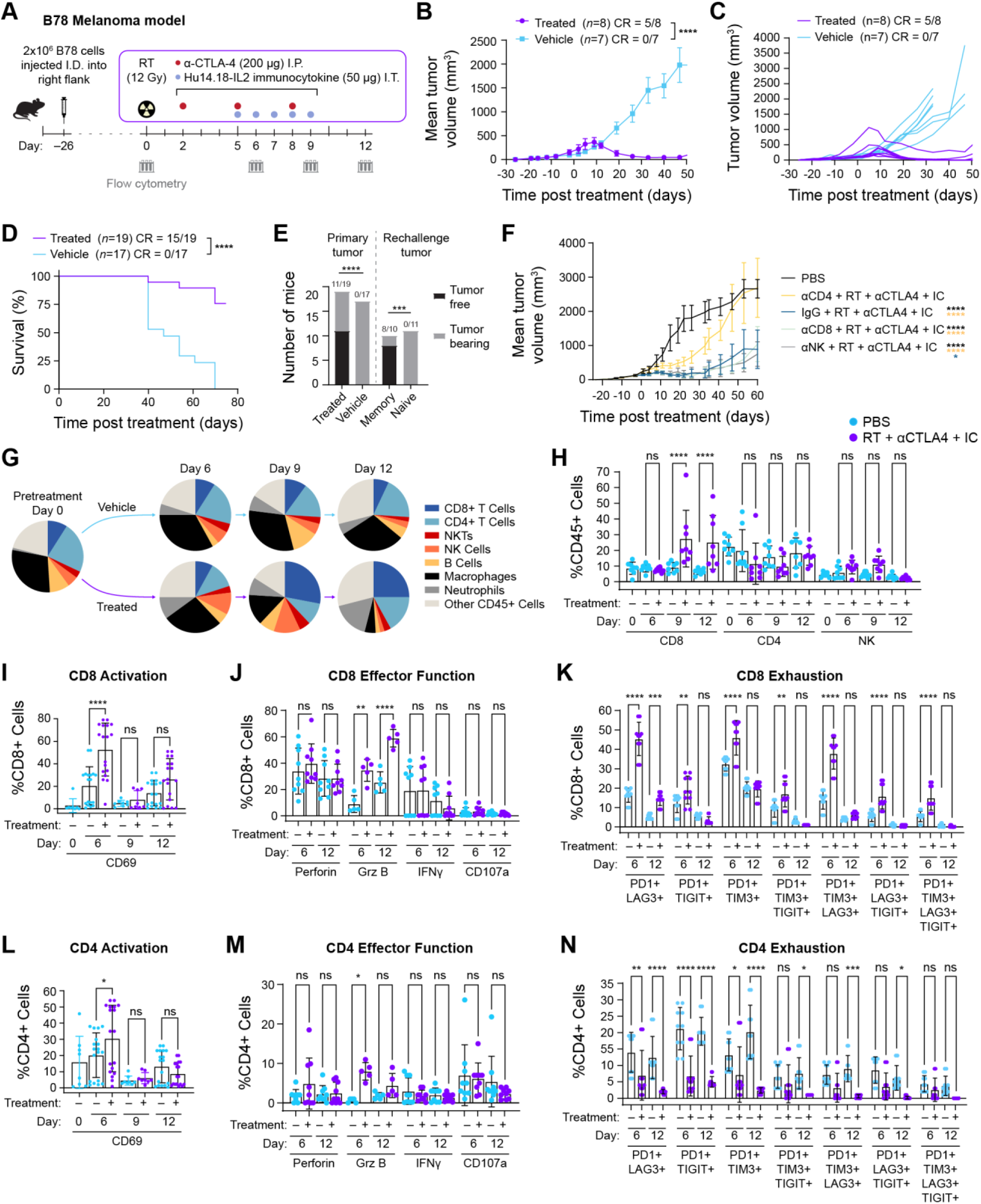
Radio-immunotherapy leads to durable responses and immune infiltrate changes in B78 melanoma model. **A)** Experimental timeline with flow cytometry. **B)** Mean tumor-growth curves for treated and vehicle B78-bearing mice (pairwise contrasts, *n*=7-8 mice/group, mean ± SEM, 1 representative experiment shown, 3 experimental repeats). **C)** Individual tumor-growth curves for treated and vehicle B78-bearing mice (*n*=7-8 mice/group, 1 representative experiment shown, 2 experimental repeats). **D)** Kaplan-Meier survival curve of treated and vehicle B78-bearing mice (pairwise log-rank test, *n*=17-19 mice/group, 3 experimental repeats combined). **E)** Number of tumor-free mice from treated and vehicle primary tumors and from treated and naïve tumor rechallenge (Fisher exact test, primary *n*=17-19, rechallenge *n*=10-11 mice/group, 3 experimental repeats). **F)** Mean tumor-growth curves during CD8, CD4, and NK cell depletion for B78-bearing mice (*n*=10 mice/group, pairwise contrasts, 1 experimental replicate, mean ± SEM). **G)** Flow cytometry of immune cell subsets in tumors from treated and vehicle B78-bearing mice across treatment time (*n*=8 mice/day, 2 experimental repeats, %CD45+ cells). **H)** Flow cytometry of CD8, CD4, NK cell changes from treated and vehicle B78-bearing mice across treatment time (*n*=10 mice/group, one-way ANOVA, 2 experimental repeats, mean ± SD). **I-K)** Flow cytometry of CD8 and **L-N)** CD4 T cell activation, effector function, and exhaustion from treated and vehicle B78-bearing mice across treatment time (*n*=8-18 mice/group, one-way ANOVA, mean ± SD, %CD8+ or %CD4+ cells, 2-4 experimental repeats).

### Cold B78 melanoma cures driven by CD4 T cells

Recently we showed that B78-bearing mice (∼50 mm^3^) that can be cured with RT and Hu14.18-IL2 alone require CD4 T cells to achieve cures and memory responses, but not NK and CD8 T cells.^19^ To probe which immune cells are driving the B78 responses and memory in these larger tumors when αCTLA-4 is included, we selectively depleted CD8, CD4, or NK cells during therapy. Treated B78-bearing mice depleted of CD8 or NK cells behaved similarly to treated mice with IgG control antibody – potent tumor-growth inhibition was seen in all 3 of these groups (**Fig 3F**) and 40-70% of the mice had complete responses. In contrast, treated B78-bearing mice depleted of CD4 cells did not clear tumors and responded no differently than mice that received PBS alone, but significantly differed from αCD8, αNK, IgG groups (**Fig. 3F**). This work highlights that CD4 cells are critical for B78 tumor response to our radio-immunotherapy *in vivo*, consistent with previous findings.^19^ We further explored immune function during radio-immunotherapy using multi-parameter flow cytometry on day 0, 6, 9, and 12. We showed that the tumor infiltrating immune populations changed over time and with therapy including a significant expansion of CD8 cells on day 9 and 12, a significant reduction in macrophages on day 9 and 12, and a significant expansion of neutrophils on day 12 (**Fig. 3G-H**, **Extended Data Fig. 3F**). Functional measurements showed that CD8 cells from treated mice are significantly activated on day 6 (**Fig. 3I**), contain significantly higher granzyme B on day 6 and 12, but express minimal to no CD107a (**Fig. 3J**). Additionally, CD8 cells from treated mice express significantly higher exhaustion markers compared to vehicle mice (**Fig. 3K**). In contrast, CD4 cells from treated mice were significantly more activated on day 6 (**Fig. 3L**), contained significantly higher granzyme B on day 6, expressed some CD107a (∼5%) though it was not different than vehicle mice (**Fig. 3M**), and expressed significantly less exhaustion markers compared to vehicle mice (**Fig. 3N**). NK cells from treated mice were significantly more activated on day 6 and 12, contained significantly higher cytolytic granules on day 6 and 12, expressed minimal to no CD107a, and expressed low levels of exhaustion markers compared to vehicle mice (**Extended Data Fig. 3G-I**). In summary, antibody depletion experiments and a multi-parameter flow cytometry time-course show CD4 T cells drive B78 tumor cures across treatment time, presumably through combination effector and helper function. CD8 T cells are not critical to tumor cures. Though CD8 T cells do become activated and express cytolytic granules, no direct killing is observed (no CD107a degranulation) and exhaustion markers are high with treatment.

### In vivo imaging captures B78 CD8 T cell atypical metabolic dynamics, with no glycolytic shift observed

To further understand why CD8 T cells are not functional in this cold B78 model, we performed *in vivo* label-free OMI to track single CD8 T cell metabolic changes across treatment on day 0, 6, 9, and 12 (**Fig. 4A**). Representative fluorescence intensity images showed decreased CD8 T cell infiltration (white outlines mark CD8 T cells) in treated mice versus vehicle mice, especially at day 6 and 9 (**Fig. 4B**, top row). Representative NAD(P)H α_1_ and FAD α_1_ images showed that CD8 T cell NAD(P)H and FAD protein-binding changes are minimal over treatment time-course (**Fig. 4B**, middle and bottom rows). Single cell CD8 T cell metabolic changes showed no significant changes in NAD(P)H α_1_ on any day, though Glass’s Δ indicated a trend towards decreased NAD(P)H α_1_ indicating an oxidative metabolism shift (**Fig. 4C**). CD8 T cell metabolic changes showed an increase in FAD α_1_ on day 9 only, indicating an oxidative metabolism shift (**Fig. 4D**). Despite the minimal metabolic changes, OMI showed that CD8 T cell size significantly increased on day 6 with an increasing trend on day 9 (**Fig. 4E**). We trained a binary classifier on CD8 T cell *in vivo* OMI variables and tested sensitivity to treated vs. vehicle condition. On day 9, the classifier identified CD8 T cells from treated versus vehicle mice with high accuracy (AUC = 0.90, **Fig. 4F**).

**Figure 4.**
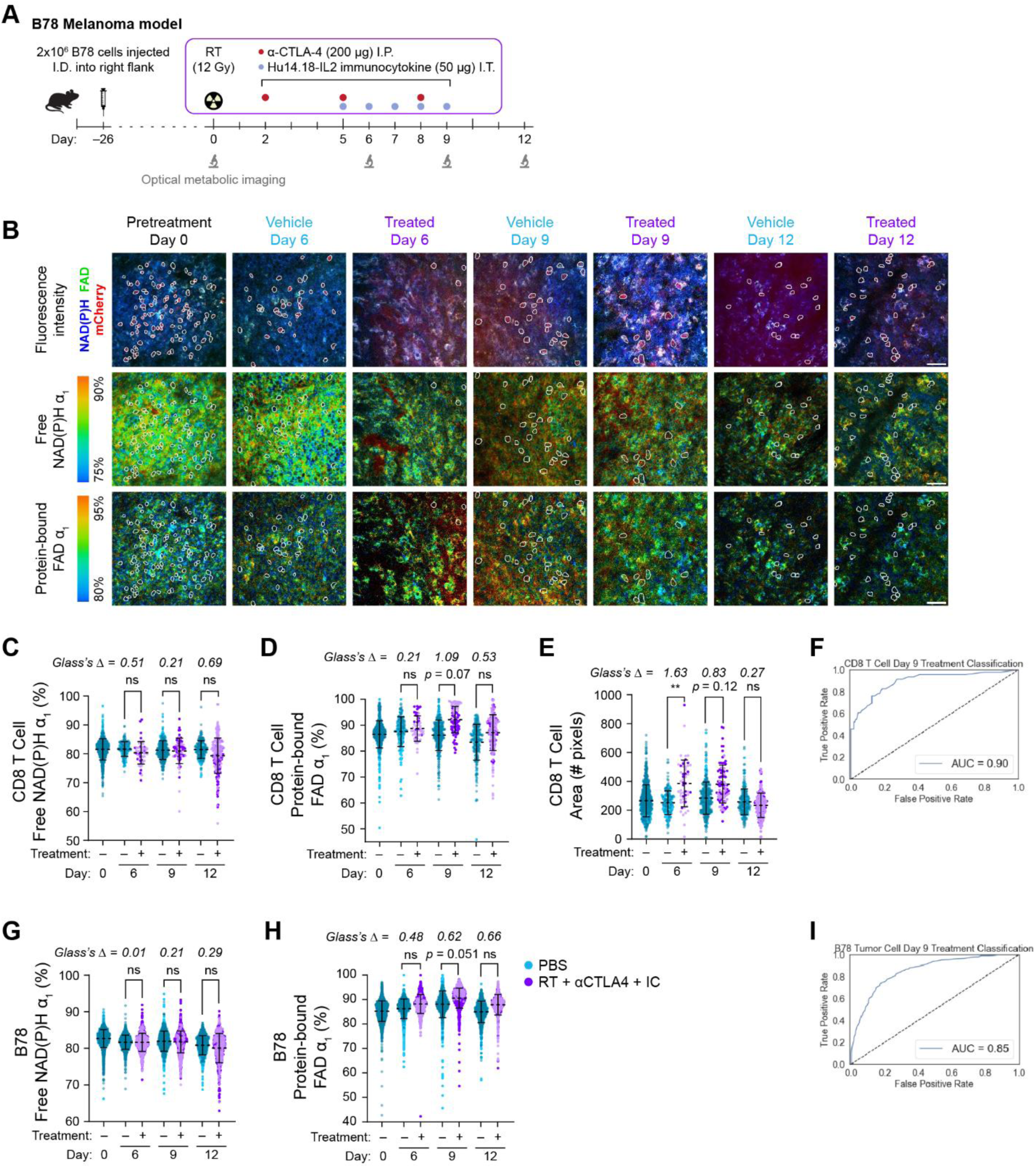
In vivo fluorescence intensity and lifetime imaging of CD8 T cells and B78 melanoma tumors shows metabolism changes with treatment. **A)** Experimental timeline with *in vivo* OMI. **B)** Representative *in vivo* B78 melanoma tumor images across time. Fluorescence intensity images (top row) show infiltrating CD8 T cells (red) expressing mCherry and autofluorescent NAD(P)H (blue) and FAD (green) present in all cells. NAD(P)H α_1_ (middle row) and FAD α_1_ (bottom row) images show spatial distribution of metabolic changes within the tumors. CD8 T cell outlines (white) overlaid on all images for clarity (1 experiment shown, 2 experimental repeats performed, scale bar 50 μm). **C-E)** CD8 T cell NAD(P)H α_1_ (proportion of free NAD(P)H), FAD α_1_ (proportion of protein-bound FAD), and area changes during therapy. **F)** ROC curve (AUC = 0.90) classifying treated or vehicle CD8 T cells from day 9. **G-H)** B78 tumor cell NAD(P)H α_1_ and FAD α_1_ changes during therapy. **I)** ROC curve (AUC = 0.85) classifying treated or vehicle B78 tumor cells from treatment day 9. **C-E)** *n*=4 mice/day, Day 0 *n*=794 cells, Day 6 *n*=247 cells, Day 9 *n*=463 cells, Day 12 *n*=438 cells, CD8 T cells imaged, mixed-effects ANOVA and Glass’s Δ, mean ± SD, 2 biological replicates. **G-H)** *n*=4 mice/day, *n*=2-4 mice/group, Day 0 *n*=2587 cells, Day 6 *n*=2521 cells, Day 9 *n*=2203 cells, Day 12 *n*=1841 cells, B78 tumor cells imaged, mixed-effects ANOVA and Glass’s Δ, mean ± SD, 2 experimental repeats. **F, I)** Random Forest algorithm, 50% train:50% test, **Table S1** shows 8 variables included, day 0 data not included.

### In vivo imaging captures B78 tumor cell metabolic dynamics during radio-immunotherapy

We also captured single B78 tumor cell metabolic changes across treatment (**Fig. 4A**). Representative fluorescence intensity images showed changes in TME over time with lower tumor cell density later in treatment (**Fig. 4B**, top row). Representative NAD(P)H α_1_ and FAD α_1_ images showed that B78 tumor cell NAD(P)H and FAD mean protein-binding minimally change over treatment time-course though heterogeneity was observed (**Fig. 4B**, middle and bottom rows). Quantified single cell B78 metabolic changes showed no changes in NAD(P)H α_1_ on any treatment day (**Fig. 4G**) and an increase in FAD α_1_ on day 9 only, indicating a shift towards oxidative metabolism (**Fig. 4H**). These metabolic shifts were validated by plating B78 *in vitro* and applying known metabolic inhibitors including NaCN and 2DG (Extended **Data Fig. 4F-I**). We trained a binary classifier on B78 tumor cell *in vivo* OMI variables and tested sensitivity to treated vs vehicle condition. On day 9, the classifier identified B78 tumor cells from treated versus vehicle mice with modest accuracy (AUC of ROC = 0.85, **Fig. 4I**).

### scRNAseq confirms B78 CD8 T cell metabolic gene expression changes with radio-immunotherapy

Next, using single-cell RNA sequencing (scRNAseq) data generated from our lab,^19^ we assessed metabolism-related gene expression changes for CD8 T cells during radio-immunotherapy to corroborate our *in vivo* OMI findings. B78-bearing C57BL/6 mice were treated with RT and Hu14.18-IL2 immunocytokine and tumors were harvested for scRNAseq on Day 8 (**Fig. 5A**).^19^ Within the CD45+ cell population, CD8, CD4, and NK cells were clustered by treatment group showing these cells were more metabolically active in treated mice with increased gene expression from several KEGG^75–77^ metabolism pathways (**Fig. 5B-C**). CD8 cells from treated mice most upregulated metabolic genes related to oxidative phosphorylation, mirroring OMI results, while CD4 cells most upregulated metabolic genes related to arginine and proline metabolism (**Fig. 5C**). Within CD8 T cells, we observed striking cellular heterogeneity differences between treated and vehicle mice including reduction of subset 1 and expansion of subsets 2-6 in treated mice (**Fig. 5D**). Subsets 5 and 6 showed the highest metabolic activity gene expression with treatment including an upregulation of genes related to pentose phosphate pathway (PPP) and glycolysis/gluconeogenesis (subset 5) and glycolysis/gluconeogenesis, PPP, oxidative phosphorylation, pyruvate metabolism, and fatty acid metabolism within subset 6 (**Fig. 5E**). These subsets expressed genes related to DNA repair and chromatin activity (*Rrm1*, *Hist1h2am*, *Hist1h2ae*, *Hist1h1b*, *Top2a*, *Pclaf*), mitosis (*Spc24*, *Cks1b*, *Smc1*), proliferation and apoptosis suppression (*Mki67*, *Birc5*), chemotaxis (*Xcl1*), and expressed cytotoxic and stem-like gene signatures (**Fig. 5F, H**). Furthermore, subsets 1 and 2 expressed significantly higher gene expression of glycolytic genes (*Gpi1, Pgm2*) while subsets 5 and 6 expressed significantly higher gene expression of oxidative genes (*Ndufa13*, *Cox6a1*) (**Fig. 5G**). To complement the metabolic characterization of the CD8 subsets, we also examined KEGG immune functional pathways and observed that subsets 5 and 6 exhibited enhanced functionality after treatment, including increased T cell receptor (TCR) signaling and IL-17 signaling signatures (subset 5) and FCε RI signaling and IL-17 signaling signatures (subset 6) **(Fig. 5I)**. In summary, gene expression changes related to metabolism and immune function indicated CD8 T cells are phenotypically changed in treated mice.

**Figure 5.**
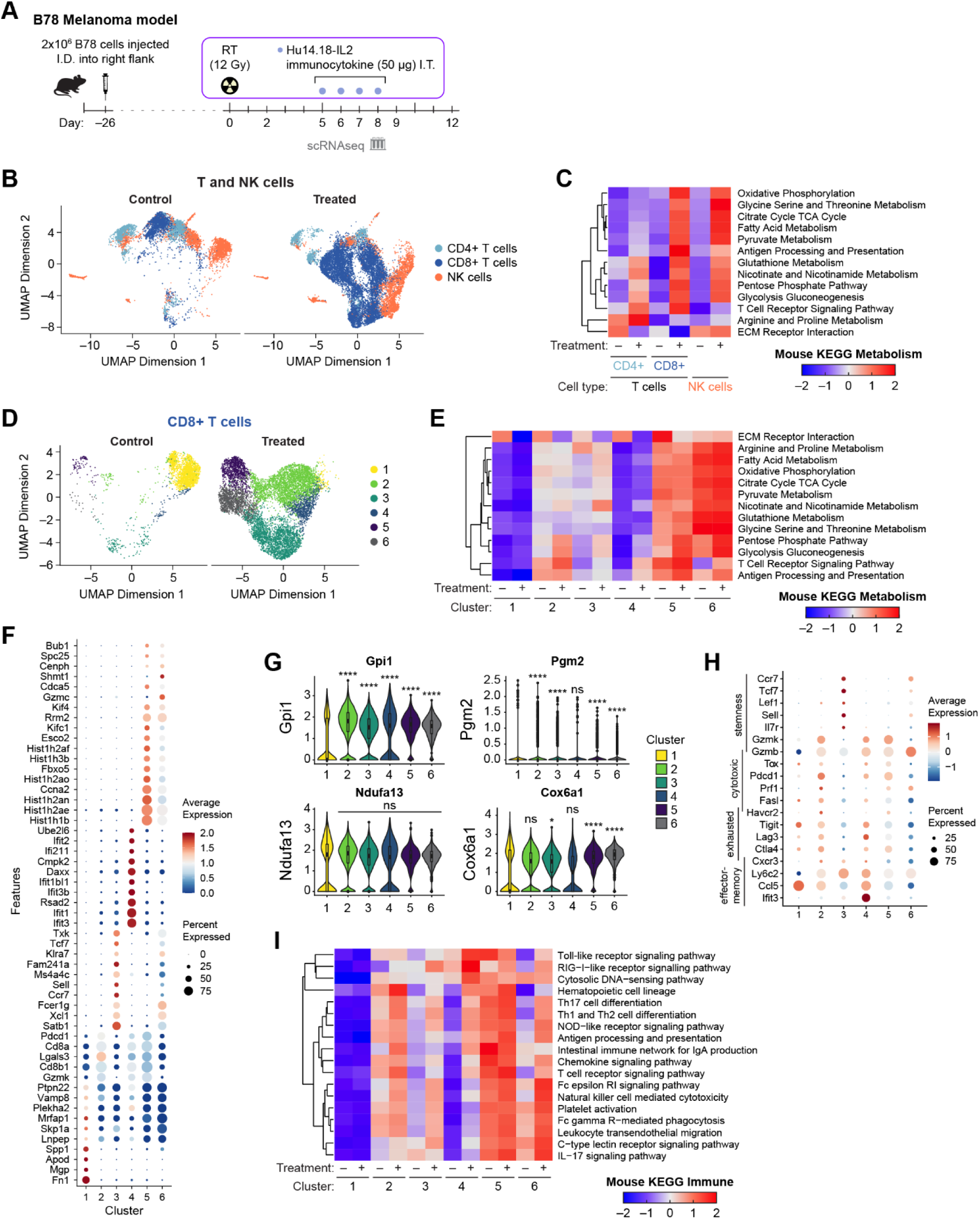
B78 immune cell scRNAseq highlights metabolic gene expression changes during radio-immunotherapy. **A)** Experimental timeline with scRNAseq. **B)** UMAP projection of T and NK cells from treated and vehicle B78-bearing mice. **C)** Heatmap of CD8, CD4, and NK KEGG metabolism average gene expression changes within treated and vehicle B78-bearing mice. **D)** UMAP projection of CD8 T cell subsets from treated and vehicle B78-bearing mice. **E)** Heatmap of CD8 KEGG metabolism average gene expression changes from treated and vehicle B78-bearing mice. **F)** Dot Plot of top 10 genes expressed by each CD8 T cell subset (6 subsets) from treated and vehicle B78-bearing mice. **G)** Quantified gene expression changes for glycolysis associated genes (*Gpi1*, *Pgm2*) and oxidative phosphorylation associated genes (*Ndufa13*, *Cox6a1*) across CD8 T cell subsets from treated and vehicle B78-bearing mice (Wilcoxon rank-sum test with Benjamini-Hochberg correction). **H)** Dot Plot of CD8 T cell phenotypes across cell subsets from treated and vehicle B78-bearing mice. **I)** Heatmap of CD8 KEGG immune function average gene expression changes within treated and vehicle B78-bearing mice. **B-I)** *n*=2 mice/treatment group, 1 experimental replicate.

### scRNAseq analysis of B78 tumors shows tumor metabolic shifts linked to radio-immunotherapy

We also reanalyzed the GD2+ B78 tumor cell populations within our scRNAseq data^19^ (**Fig. 6A**) and observed that *Sox10* expressing tumor cells from vehicle and treated mice differed when two major tumor clusters were selected (**Fig. 6B**). Tumors from vehicle mice expressed upregulated genes related to cell proliferation/motility/invasion (*Gpc3*, *Tspan13*, *Ccpg1*, *Asah1, Itfg1*), metabolism (*Cyp20a1*, *Aldoc*), enzymes that degrade gangliosides (*Hexb*), inhibition of apoptosis and promotion of angiogenesis (*Srpx2*, *Capn6*), promotion of fibroblasts and collagen deposition (*Glg1*, *Fmod*), and others (subset 0) (**Fig. 6C**). On the other hand, tumors from treated mice upregulated genes related to both inhibition (*Ifit3*, *Ifit3b*) and promotion (*Xaf1*) of apoptosis, increased PD-L1 expression (*Cd274*), increased GTPase activity tied to cell signaling/proliferation/migration/cytoskeletal organization (*Gbp6*, *Tgtp1*, *Irgm1*, *Iigp1*, *Igtp*, *Gbp3*, *Gbp2*), and interferon signaling (*Ifi47*, *Apol9b*) (subset 1) (**Fig. 6C**). KEGG metabolism pathway signatures are distinct between the two subsets, where subset 0 showed a downregulation of metabolic genes and activity in the treated mice versus high arginine/proline metabolism within the vehicle mice (**Fig. 6D**). Subset 1 from treated mice also showed a downregulation of most metabolic genes and activity, except for increased nicotinate/nicotinamide metabolism, while subset 1 from vehicle mice expressed an increase in genes related to glycine/serine/threonine metabolism, glutathione metabolism, and citrate cycle/TCA cycle (**Fig. 6D**). Subset 0 also expressed significantly higher gene expression of glycolytic genes (*Gpi1, Pgam1*) and while subset 1 expressed significantly higher gene expression of one of the oxidative genes (*Sdha*) (**Fig. 6E**). We also observed an upregulation in genes associated with apoptosis within tumor subset 1 (**Fig. 6F**). In summary, we observed that B78 tumor cells from vehicle mice versus treated mice have distinct gene expression in metabolism and related functions based on scRNAseq.

**Figure 6.**
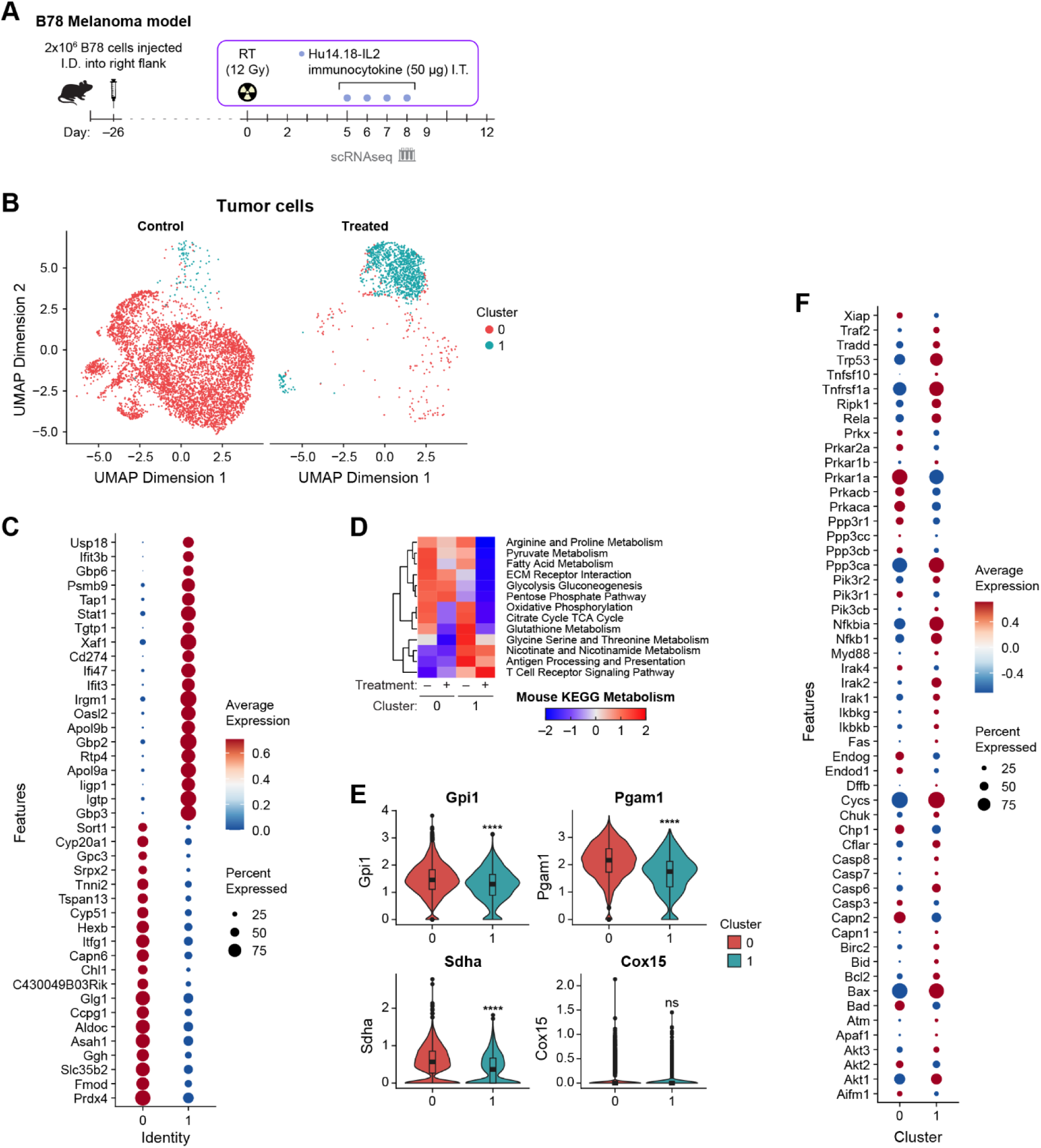
B78 tumor cell scRNAseq highlights tumor metabolic gene expression changes during radio-immunotherapy. **A)** Experimental timeline with scRNAseq. **B)** UMAP projection of B78 tumor cells from treated and vehicle B78-bearing mice. **C)** Dot Plot of top 20 genes expressed by the two B78 tumor subsets. **D)** Heatmap of B78 KEGG metabolism average gene expression changes from treated and vehicle B78-bearing mice. **E)** Quantified gene expression changes for glycolysis associated genes (*Gpi1*, *Pgm2*) and oxidative phosphorylation associated genes (*Sdha*, *Cox15*) across tumor cell subsets from treated and vehicle B78-bearing mice (Wilcoxon rank-sum test with Benjamini-Hochberg correction). **F)** Dot Plot of top 40 KEGG apoptosis genes for each B78 tumor cell subset from treated and vehicle B78-bearing mice. **B-F)** *n*=2 mice/treatment group, 1 experimental replicate.

### Multiplex immunofluorescence spatial profiling of B78 tumor and lymph node elucidates infiltration changes and immune-tumor interactions with radio-immunotherapy

Finally, we performed multiplex immunofluorescence on B78 tumors and lymph nodes on day 0, 6, 9, and 12 (**Fig. 7A**). Tumors were stained nuclei/DNA (DAPI), immune cells (CD4, CD8α, CD161), tumor cells (SOX10), cytolytic granules (perforin), and cell death (cPARP-1) (**Fig. 7B, Extended Data Fig. 6A**). Representative tumor regions of interest (ROI) across treatment time showed changes in immune infiltration, tumor organization, and cell death (**Fig. 7B, Extended Data Fig. 6A**). Tumor ROI from treated mice showed significantly increased CD8 and CD4 infiltration and significantly decreased tumor cells on Day 12 (**Fig. 7C-E**), while perforin significantly increased on Day 6 (**Extended Data Fig. 6B**). Spatial profiling via distance ratio from CD4 cells to nearest SOX10+ tumor cell revealed that CD4 cells were originally farther away in treated mice on Day 6 but then transitioned to being closer to tumor cells on Days 9 and 12, compared to vehicle mice (**Fig. 7F**). In contrast, the distance ratio from CD8 cells to nearest SOX10+ tumor cell showed no difference with treatment on Day 6, but CD8 cells from treated mice transitioned closer to tumor cells on Days 9 and 12 (**Fig. 7G**). Tumor draining lymph nodes (TDLN) were stained for nuclei/DNA (DAPI), immune cells (CD3, CD4, CD8α, CD19, F4/80), and cell activation (CD69) (**Fig. 7H, Extended Data Fig. 6D-E**). Lymph nodes from treated mice showed increased numbers of B cells plus CD4 and CD8 cells on Days 9 (peak) and 12 compared to vehicle mice (**Fig. 7I-K, Extended Data Fig. 5A,C**), suggesting immune priming and activation is more robust in treated mouse lymph nodes. This was also observed in TDLN of mice treated with RT and Hu14.18-IL2 alone.^19^

**Figure 7.**
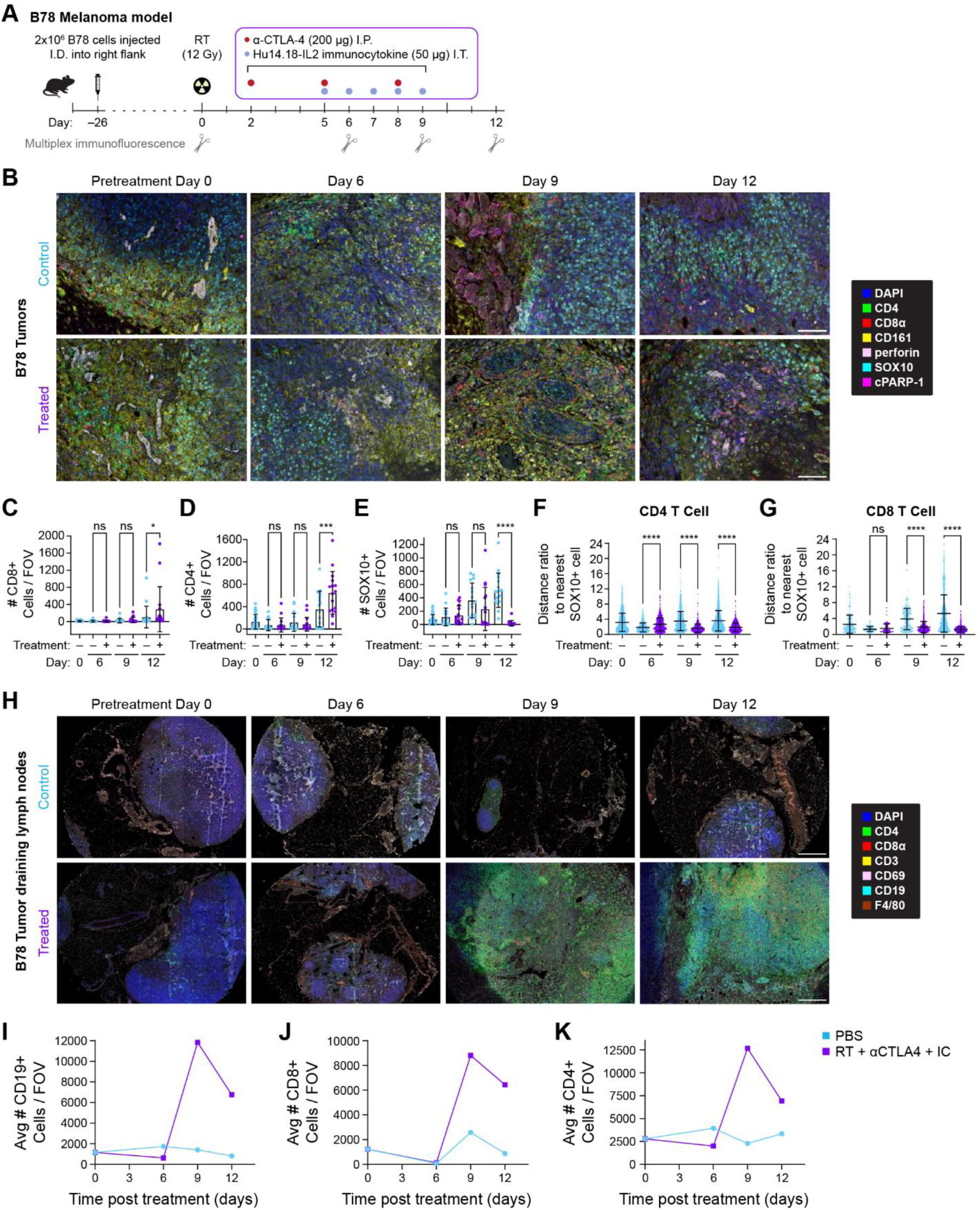
Multiplex immunofluorescence spatial profiling of B78 tumor and lymph node tissues elucidates immune-tumor interactions and infiltration changes with radio-immunotherapy. **A)** Experimental timeline with multiplex immunofluorescence. **B)** 20× mIF images from B78 tumors across treatment time-course from vehicle (top row) and treated mice (bottom row) (*n*=2-4 mice/group, 1 experimental replicate, scale bar 100 μm). **C-E)** Quantified CD8+, CD4+, and SOX10+ tumor cells from treated and vehicle B78-bearing mice across treatment time. **F-G)** CD4 and CD8 T cell distance ratio from nearest tumor cell using nearest-neighbor. **H)** mIF images from B78 tumor-draining lymph nodes across treatment time-course from vehicle (top row) and treated mice (bottom row). 3×3 20× images stitched together/lymph node (*n*=1-2 mice/group, *n*=9 images/lymph node, 1 experimental replicate, scale bar 300 μm). **I-K)** Quantified CD19+, CD8+, and CD4+ immune cells from lymph nodes from treated and vehicle B78-bearing mice across treatment time (*n*=1-2 mice/group, *n*=2-4 lymph node/group, 1 experimental replicate, average shown). **C-G)** *n*=2-4 mice/group, *n*=8 images/tumor, one-way ANOVA, 1 experimental replicate, mean ± SD.

### DISCUSSION

Here, we executed a multimodal approach to investigate the therapeutic effects and mechanism of our curative radio-immunotherapy regimen on two contrasting tumor models: immunologically-hot MC38 colon carcinoma and immunologically-cold B78 melanoma. We created an immunocompetent mouse model with CD8 T cells that express mCherry to easily track and monitor CD8 T cell dynamics *in vivo*.

The well-studied MC38 tumor model is minimally sensitive to some monotherapies such as glutamine antagonists,^49^ and very sensitive to combination therapies such as αPD-1/αCTLA-4 blockade (75% tumor free mice)^78^ or PD-1/glutamine antagonist (90% tumor free mice).^49^ Here, we showed that MC38 tumors were highly sensitive to our radio-immunotherapy combination with nearly 100% primary and rechallenged tumor rejection. These tumor responses were primarily mediated by CD8 T cells, as shown by others^73,78,79^, though NK and CD4 T cells were also engaged. CD8 T cell activation took place early in therapy (day 6 versus day 9) based on CD69 expression and increased cell size measured by *in vivo* OMI. As expected, metabolic imaging showed that CD8 T cells in the MC38 TME increased glycolysis after priming/activation, fueling their effector function (day 6).^80,81^ This is mirrored by increased cytolytic granules and active degranulation shown also on day 6 via flow cytometry. Later (day 9) imaging showed that this immune response was dampened with CD8 T cells shifting their metabolism back towards an oxidative phenotype, consistent with transition to a memory-like state as others reported.^49,79,82^ We also showed that this oxidative phenotype is not due to exhaustion. These MC38-TME metabolic dynamics are consistent with traditional assays, like Seahorse analysis,^49,82^ though this is the first time these changes have been shown via *in vivo* label-free metabolic imaging. Interestingly, when we trained a machine learning classifier on CD8 T or MC38 tumor cells *in vivo* metabolic variables, treated versus vehicle mice were accurately classified – showing that our radio-immunotherapy prompts unique metabolic signatures in both CD8 T and MC38 cells.

In contrast, the cold GD2+, no MHC-I B78 model is difficult to treat, does not respond to ICI alone, and requires combination therapy for tumor response.^10,17,18,73^ B78 tumors do respond to the combination of radiation, αCTLA-4, and Hu14.18-IL2 immunocytokine^16,17^ with tumor rejections in 58% of primary tumors and 80% of rechallenged tumors. Interestingly, our immune depletion studies showed that these tumor responses are primarily driven by a CD4 T cell immune response, which has been shown previously,^18,19,83,84^ though CD8 T and NK cells were also activated and contained cytolytic granules. Treatment did not substantially increase CD4 infiltration into the TME but caused a large influx of CD4 T cells into the tumor draining lymph nodes, suggesting a key role of CD4 T cells in immune priming, cell signaling, and helper function. A detailed analysis of CD4 function and mechanism within B78 is explored by Erbe *et al.,* which shows that B78 cells upregulate some MHC-II *in vivo*, but not MHC-I, and can be directly killed by CD4 cells.^19,84^ Here we focus on the different roles of CD8 T cells in B78 vs. MC38. Flow cytometry and *in vivo* imaging showed that CD8 T cell activation took place early in therapy for B78 (day 6), based on CD69 expression and increased cell size. Surprisingly, this CD8 T cell priming/activation was not accompanied by the expected metabolic shift towards glycolysis^85^ as seen in MC38; instead the treated CD8 T cells in the B78-bearing mice showed no difference in metabolic profile compared to vehicle. Functionally, the treated CD8 T cells contained increased granzyme B on day 6 and 12 but there was no evidence of cytolytic activity on either day, unlike the CD8 cytolytic activity observed in MC38. As treatment progressed, CD8 T cells exhibited an oxidative shift in their metabolism, consistent with a memory or exhausted phenotype, as shown by *in vivo* imaging and scRNAseq. Flow cytometry confirmed increased expression of multiple exhaustion markers on CD8 T cells but not CD4 T cells from B78 treated mice. This lack of effector function, together with the exhaustion and oxidative phenotype, is partly due to the minimal to no MHC-I expressed by B78 melanoma tumors, which recapitulates a common occurrence in the clinical presentation of melanoma.^38,86–89^ CD8 T cells are primed and activated early but never receive the full stimulatory signal required to upregulate glycolytic metabolism. Additionally, as little to no MHC-I is present on the tumor cells, these CD8 T cells are not mediating tumor killing, but experience chronic stimulation, likely from cytokines released by the activated CD4 and NK cells, which may lead to exhaustion. Gene expression changes showed that the subsets of CD8 T cells most expanded in treated mice (subsets 5 and 6) upregulated genes related to DNA repair and chromatin activity, mitosis, chemotaxis, and suppression of apoptosis as well as functional changes including increased TCR and IL-17 signaling. Phenotypically, treated CD8 T cell subsets were diverse with gene expression related to stem-like, cytotoxic, exhausted, and effector memory phenotypes. When we trained a machine learning classifier on B78 CD8 T cell *in vivo* metabolic variables, treated mice versus vehicle mice were consistently classified – showing that our radio-immunotherapy prompts unique metabolic signatures even when CD8 T cells are not driving tumor response. Others have imaged metabolism in immune cells *in vitro* or *in vivo* in a bulk context during immunotherapy^90–92^ but, to the best of our knowledge, this is the first time metabolic imaging has monitored changes induced by curative immunotherapy at the single-cell level *in vivo*.

B78 tumor cells showed similar metabolic changes to infiltrating CD8 T cells with radio-immunotherapy, where treated tumor cells shifted to a more oxidative metabolism over time, especially on day 9 and 12, via OMI. Gene expression changes of KEGG metabolism pathways showed that tumor cells from treated mice were less metabolically active compared to vehicle mice, indicating that most tumor cells were dying. The remaining tumor cells within treated mice upregulated gene expression related to PD-L1, apoptosis, and increased proliferation/migration. Interestingly, the tumor cells from vehicle mice upregulated gene expression to promote tumor progression including motility, angiogenesis, fibroblast and collagen interactions, and downregulation of apoptosis. Our radio-immunotherapy reduces tumor infiltrating macrophages (involved in collagen production and organization) and promotes a healthy collagen phenotype – which may contribute to the change in gene expression observed within treated mice.^36,93^ We also observed in the treated mice an increase in genes associated with programmed cell death, NF-κB signaling, and p53 tumor suppressor activity that imply a complex combination of both pro- and anti-apoptotic function within the treated mice.^94–96^ Despite these complex mechanisms of tumor resistance and escape (PD-L1 increase, migratory phenotype, apoptosis genes), 58% of mice became tumor free after radio-immunotherapy to the primary tumor. Both CD4 and CD8 T cells from treated mice were also in closer proximity to tumor cells compared to vehicle mice, which has been correlated to improved patient survival.^97–100^ For the CD4 cells this would be consistent with immune-tumor interaction and immune mediated killing of the MHC-II expressing B78 cells.^84^ For the CD8 cells, increased proximity to B78 tumor cells that do not express any detectible MHC-I may be an indicator that they were actually closer to antigen presenting cells (APCs) located near tumor cells. Perhaps tumor peptides were presented by APC MHC-I to enable CD8 activation and possibly killing of APCs.

Overall, we have shown that curative radio-immunotherapy across hot versus cold tumors produces divergent CD8 T cell metabolic, functional, and genetic phenotypes that correlate to their distinct roles in tumor immunity. Had we only investigated CD8 T cell activation, we would have missed these complex and divergent phenotypes. This work suggests the importance of developing combination therapies that target CD8 T cell exhaustion and highlights the key role that other immune cells (CD4 T, NK) play during tumor response. We believe this is the first time label-free *in vivo* metabolic imaging has been used to study single-cell effects of radio-immunotherapy in preclinical models and correlate the imaging findings at a deep molecular level to multi-parameter flow cytometry, scRNAseq, and mIF.

## METHODS

### Cell lines

B78-D14 (B78) melanoma is a poorly immunogenic mouse cell line derived from B78-H1 melanoma cells, which were originally derived from B16 melanoma. These cells were obtained from Ralph Reisfeld (Scripps Research Institute) in 2002. B78 cells were transfected with functional GD2/GD3 synthase to express the disialoganglioside GD2, which is overexpressed on the surface of many human tumors including melanoma. These B78 cells were also found to lack melanin. MC38 is a mouse colon adenocarcinoma cell line. These cells were obtained from Brad St. Croix (NIH). B78 and MC38 cells were grown in RPMI-1640 (Gibco) supplemented with 10% FBS and 1% penicillin/ streptomycin; B78 media was periodically supplemented with 400 μg G418 and 500 μg Hygromycin B/mL. Mycoplasma testing was performed every 6 months.

### Animal models

All animal work was performed under an animal protocol approved by the Institutional Animal Care and Use Committee at the University of Wisconsin, Madison. A maximum of 5 mice were housed/cage. Standard chow and water were provided *ad libitum*. All mice were housed with a 12 h/12 h dark/light cycle at 22–23 °C and 40–50% humidity. Enrichment (nesting material and igloo) was provided to all cages. Male and female mice were used for all experiments at 8-24 weeks of age. C57BL/6J mice were obtained from Jackson Laboratories. C57BL/6NTac were obtained from Taconic Biosciences. The mCherry-CD8α knock-in mouse (CD8-mCherry) was designed and generated by the University of Wisconsin-Madison Genome Editing (RRID:SCR_021070) and Animal Model (RRID:SCR_024797) Cores. Pronuclear microinjection was conducted into C57BL/6J embryos using a mixture of Cas9 protein, a long single-stranded donor template (IDT, Megamer ssDNA Fragment), and an *in vitro* transcribed sgRNA (TGCTGGTGGAGAGCACACCATGG) targeting the N-terminus of the CD8α coding sequence. The mCherry-P2A knock-in cassette resulted in a CD8 reporter allele, rather than an mCherry-CD8 fusion. Pups were sequenced using primers 1033 and 1185 (N-terminal) and 1034 and 800 (C-terminal) followed by Sanger sequencing. A sequence-confirmed founder was identified, the founder was backcrossed to C57BL/6J to produce F1 progeny, and F1 progeny were similarly sequence confirmed.

### Mouse tumor model preparation and therapy administration

2×10^6^ B78 cells were injected intradermally (I.D.) into the right flank in a volume of 0.1 mL PBS. 5×10^5^ MC38 cells were injected I.D. into the right flank in a volume of 0.1 mL PBS. For the GD2+ B78 model, mice were randomized into treatment groups when tumors reached enrollment size (∼150 mm^3^), which typically required 3 to 4 weeks of *in vivo* growth. For the GD2-MC38 model, mice were randomized into treatment groups when tumors reached enrollment size (∼150 mm^3^), which typically required 7 to 10 days of *in vivo* growth. Each randomized group included male and female mice, where the average tumor size of each group was matched. The first day of treatment is defined here as “day 0”. For both B78 and MC38 mice, external beam radiation therapy was delivered to the tumor surface of treated mice only on day 0, using an X-RAD 320 cabinet irradiator system (Precision X-Ray, North Branford, Connecticut, United States). Mice were immobilized using custom lead jigs that exposed only the dorsal right flank. Radiation was delivered in one fraction to a maximum dose of 12 Gray (Gy). For both B78 and MC38 mice, systemic mouse α-CTLA-4 antibody was administered to treated mice once daily on days 2, 5, and 8 via intraperitoneal injections of 200 μg in 200 μL PBS (Bristol-Meyers Squibb, Redwood City, California, United States and NeoClone Biotechnologies International, Madison, Wisconsin, United States). For B78 mice only, Hu14.18-IL2 immunocytokine, a monoclonal anti-GD2 antibody fused to IL-2 cytokine, was administered to treated mice once daily on days 5 to 9 via intratumoral injections of 50 μg in 100 μL PBS (AnYxis Immuno-Oncology GmbH, Vienna, Austria). For MC38 mice only, IL-2 cytokine was administered to treated mice once daily on days 5 to 9 via intratumoral injections of 75,000 IU in 100 μL PBS (Hoffman-La Roche, Basel, Switzerland). Vehicle-treated mice were injected once daily on days 2, 5, and 8 via intraperitoneal injections of 200 μL PBS and once daily on days 5 to 9 via intratumoral injections of 100 μL PBS. No external beam radiation therapy was administered to vehicle-treated mice. Pretreatment mice received no PBS or immunotherapy injections and no external beam radiation therapy. Tumor-growth was monitored via caliper measurements. Tumor volumes were calculated as such: (width^2^ × length) / 2. All tumor efficacy and survival experiments in both B78 and MC38 models were completed in CD8-mCherry and C57BL/6J mice. Three experimental repeats were completed for both B78 and MC38 tumor models.

### Mouse rechallenge studies

At least two months after treated mice were cured, tumor rechallenge studies were performed. For the B78 model, 2×10^6^ B78 cells were injected I.D. into the right abdomen of both cured / memory mice and age-matched naïve mice. For the MC38 model, 5×10^5^ MC38 cells were injected I.D. into the right abdomen of both cured / memory mice and age-matched naïve mice. No treatment was given to either cured / memory mice or naïve mice. Tumor-growth was monitored via caliper measurements. Tumor volumes were calculated as such: (width^2^ × length) / 2. All rechallenge experiments in both B78 and MC38 models were completed in CD8-mCherry and C57BL/6J mice. Three experimental repeats were completed for both B78 and MC38 tumor models.

### Mouse immune depletion studies

For the B78 model, C57BL/6J mice were used for immune depletion studies. CD8 T cells were depleted via anti-CD8 antibody (clone 2.43, BioXcell) at 50 μg in 0.5 mL PBS/dose on days -2, 2, 5, 8, 12, 15, 18, 22. CD4 T cells were depleted via anti-CD4 antibody (clone GK1.5, BioXcell) at 200 μg in 0.5 mL PBS/dose on days -2, 5, 12, 18. NK cells were depleted via anti-NK1.1 antibody (clone PK136, BioXcell) 200 μg in 0.5 mL PBS/dose on days -2, 2, 5, 8, 12, 15, 18, 22. Control mice received normal rat IgG antibody (Sigma-Aldrich) 200 μg in 0.5 mL PBS/dose on days -2, 2, 5, 8, 12, 15, 18, 22. Flow cytometry of peripheral blood was performed to confirm antibody depleted was successful. For the MC38 model, C57BL/6NTac mice were used for immune depletion studies. CD8 T cells were depleted via anti-CD8 antibody (BioXcell) at 50 μg in 0.5 mL PBS/dose on days -1, 6, 13, 20. CD4 T cells were depleted via anti-CD4 antibody (BioXcell) at 200 μg in 0.5 mL PBS/dose on days -1, 6, 13, 20. NK cells were depleted via anti-NK1.1 antibody (BioXcell) 200 μg in 0.5 mL PBS/dose on days -1, 6, 13, 20. One experimental replicate was completed for B78 model and two biological replicates were completed for MC38 tumor model. Statistical comparison of different immune depleted mouse groups is color-coded for each comparison (**Figure 1, 3**).

### Multiphoton imaging in vitro

*In vitro* imaging of CD3+ T cells was performed by negative isolation of splenic T cells (STEMCELL Technologies) from a C57BL/6J mouse and CD8-mCherry reporter mouse using an EasySep Mouse T Cell Isolation Kit. Isolated T cells were plated in a PBS droplet on coated glass imaging dishes, allowed to settle for 2 hours, and then imaged the same day. Autofluorescence images were captured with a custom-built multi-photon microscope (Bruker) using an ultrafast femtosecond laser (InSight DSC, Spectra Physics). Autofluorescence images were captured with a custom-built multi-photon microscope (Bruker) using an ultrafast femtosecond laser (InSight DSC, Spectra Physics). Fluorescence lifetime measurements were performed using time-correlated single photon counting electronics (Becker & Hickl). Fluorescence emission, in FLIM mode, was detected simultaneously in three channels using bandpass filters of 466/40 nm (NAD(P)H), 514/30 nm (FAD), and 650/45 nm (mCherry) prior to detection with three GaAsP photomultiplier tubes (Hamamatsu). All three fluorophores were simultaneously excited using a previously reported wavelength mixing approach.^59,109,110^ During wavelength mixing in FLIM mode, typical power at the sample was ∼0.9-2.1 mW for the 750 nm laser and ∼3.3-5.4 mW for the 1041 laser. All images were acquired with a 40×/1.13 NA water-immersion objective (Nikon) at 512×512 pixel resolution and an optical zoom of 1.0-2.0. NAD(P)H, FAD, and mCherry intensity and lifetime images were acquired across 6 fields of view. One biological replicate was performed.

*In vitro* imaging of B78 and MC38 tumor cells was performed by plating 100,000-150,000 tumor cells in RPMI-1640 media, on coated glass imaging dishes (2 technical replicates) 48 hours prior to imaging. To inhibit oxidative phosphorylation / electron transport chain complex IV, cells were first imaged untreated, then treated with 4 mM sodium cyanide (NaCN, Thermo Scientific) in PBS, then imaged again 15 minutes post-treatment. To inhibit glycolysis, cells were first imaged untreated, then treated with 10 mM 2-Deoxy-D-glucose (2DG, Sigma-Aldrich) in PBS, then imaged again 2 hours post-treatment. B78 and MC38 cells were imaged for two experimental repeats for each inhibitor treatment. Autofluorescence images were captured with a custom-built multi-photon microscope (Bruker) using an ultrafast femtosecond laser (InSight DSC, Spectra Physics). The laser was tuned to 750 nm for NAD(P)H excitation and tuned to 890 nm for FAD excitation. The average power at the sample was ∼6-9 mW for NAD(P)H excitation and ∼10-12 mW for FAD excitation. A pixel dwell time of 4.8 μs was used for all images. NAD(P)H and FAD images were acquired sequentially. A 440/80 nm bandpass filter isolated NAD(P)H emission onto the photomultiplier tube (PMT) detector. A dichroic mirror directed wavelengths greater than 500 nm onto a 550/100 nm bandpass filter, isolating FAD emission onto a second PMT. Fluorescence lifetime measurements were acquired with time-correlated single photon counting electronics (Becker and Hickl) and a GaAsP PMT (Hamamatsu). All images were acquired with a 40×/1.13 NA water-immersion objective (Nikon) at 512×512 pixel resolution and an optical zoom of 1.0. NAD(P)H and FAD intensity and lifetime images were acquired to sample metabolic behavior of B78 and MC38 tumor cells across 4-5 fields of view/dish/condition. Two experimental repeats were performed for both B78 and MC38.

### Multiphoton imaging in vivo

All imaging experiments were completed in CD8-mCherry mice. Intravital imaging of mouse B78 tumors was performed in pretreatment mice (*n* = 4), vehicle-treated mice (*n* = 6), and treated mice (*n* = 6) across days 0, 6, 9, and 12 of therapy. Intravital imaging of mouse MC38 tumors was performed on pretreatment mice (*n* = 4), vehicle-treated mice (*n* = 4), and treated mice (*n* = 4) across days 0, 6, and 9 of therapy. Imaged mice for all work were 50% male and 50% female. Day 0 images, the pretreatment group, are used as a baseline comparison for both vehicle and radioimmunotherapy-treated groups. Immediately prior to tumor imaging, skin flap surgery exposed flank tumors as previously reported.^60,93,111–113^ Mice were anesthetized with isoflurane, and then, the skin around the tumor was cut into a flap and separated from the body cavity so that the tumor laid flat on the imaging stage while still connected to the vasculature. Mice were placed on a specialized microscope stage for imaging and kept in a heating chamber (air maintained at 37°C) during imaging. An imaging dish insert and PBS for coupling were used with surgical tape to secure skin flap tumors. All imaging experiments were terminal, with new mice being imaged at each time point. The skin flap method allowed easy tumor access for intratumoral immunotherapy administration on days 5 to 9 and eliminated the concern of an immune response related to the flank window method.

Autofluorescence images were captured with a custom-built multi-photon microscope (Bruker) using an ultrafast femtosecond laser (InSight DSC, Spectra Physics). Fluorescence lifetime measurements were performed using time-correlated single photon counting electronics (Becker & Hickl). Fluorescence emission, in FLIM mode, was detected simultaneously in three channels using bandpass filters of 466/40 nm (NAD(P)H), 514/30 nm (FAD), and 650/45 nm (mCherry) prior to detection with three GaAsP photomultiplier tubes (Hamamatsu). All three fluorophores were simultaneously excited using a previously reported wavelength mixing approach.^59,109,110^ During wavelength mixing in FLIM mode, typical power at the sample was ∼2.4-6.6 mW for the 750 nm laser and ∼4.5-9.6 mW for the 1041 laser. All images were acquired with a 40×/1.13 NA water-immersion objective (Nikon) at 512×512 pixel resolution and an optical zoom of 1.0-2.0. NAD(P)H, FAD, and mCherry intensity and lifetime images were acquired to sample metabolic behavior of B78 and MC38 tumor as well as CD8 T cells across 5-11 fields of view and multiple depths within each tumor. Within **Figure 2B, 4B** the *in vivo* multiphoton images shown in top, middle, and bottom rows are the identical fields of view represented in three different ways: intensity, free NAD(P)H α_1_, and protein-bound FAD α_1_. Two experimental repeats were performed for both B78 and MC38.

### Multiphoton image analysis

The fluorescence lifetimes of free and protein-bound NAD(P)H and FAD are distinct, and these lifetimes along with their weights can be recovered with a two-exponential fit function. Therefore, fluorescence lifetime decays for both NAD(P)H and FAD were fit to the following bi-exponential function in SPCImage (v8.1):

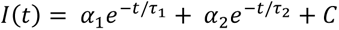

For NAD(P)H, τ_1_ corresponds to the free lifetime, τ_2_ corresponds to the protein-bound lifetime, and the weights (α_1_, α_2_; α_1_ + α_2_ = 1) correspond to the proportion of free and protein-bound NAD(P)H, respectively.^61,62,64,65,114^ Conversely for FAD, τ_1_ corresponds to the protein-bound lifetime and τ_2_ corresponds to the free lifetime.^65,67,115^ An instrument response function was measured using SHG (900 nm excitation) from urea crystals for input into the decay fit procedure. The following fluorescence lifetime endpoints were calculated from the fitted model: τ_1_, τ_2_, α_1_, and α_2_ for both NAD(P)H and FAD; along with the fluorescence lifetime redox ratio (FLIRR): NAD(P)H α_2_ / FAD α_1_ (**Extended Data Table** 1).^61,62,66,67,114,116–118^

Automated cell segmentation was performed through Cellpose2.0, with manual corrections made as needed using the Napari viewer through a custom Python script.^103,119^ Single tumor cells were identified, circled, and segmented as whole cells (nuclei and cytoplasm captured) using NAD(P)H intensity images. Single CD8 T cells were identified, circled, and segmented as whole cells using mCherry intensity images. The resulting segmented images were saved as masks. Using these masks and the raw imaging data, fluorescence lifetime variables and FLIRR values were calculated for each individual cell using custom code adapted from Cell Analysis Tools.^120^ Fluorescence lifetime variables were calculated on a region of interest or cell level, with a single decay curve for each individual cell.^121^ Receiver operator characteristic (ROC) curves were generated using a random forest algorithm. These ROC curves classified CD8 T cell and tumor cell population based on treatment conditions. **Extended Data Table 1** shows metabolic and morphological parameters included in the algorithm; 50% train: 50% test. Calculations were performed using Python (v3.10.12) / Spyder.

### Flow cytometry sample preparation and acquisition

All flow cytometry experiments were completed in CD8-mCherry mice. At each treatment timepoint, mice were euthanized with CO_2_, and tumors and spleens were excised. Spleens were manually disaggregated and filtered with a 70 μm filter (Corning), resulting in a single cell suspension. Red blood cell lysis was performed on spleens with ACK lysis buffer (Gibco) or water. Tumors were first manually disaggregated in petri dishes by chopping into ∼1 mm pieces using a scalpel. Tumors were transferred to 50 mL conical tubes containing 2.5-5 mL of complete RPMI (Gibco), 2.5 mg of collagenase type IV, 250 μg of DNase, and 1X protein transport inhibitor (PTI, eBioscience). Samples were further disaggregated by placing on a heated shaker at 37°C for 45 minutes or placed on a gentleMACS Octo Dissociator using program 37C_m_TDK1. Final single cell suspensions of tumor samples were obtained by passing through a 70 μm filter. Tumor and spleen samples were resuspended in flow buffer (PBS + 2% fetal bovine serum) plus 1X PTI, kept on ice, and protected from light for 4-6 hours. 3×10^6^ cells from each individual tumor or spleen sample were added to individual 5 mL flow tubes. Leftover cells were pooled and used for fluorescence minus one controls. For the live/dead control, 1.5×10^6^ cells were killed via heat shock and mixed with 1.5×10^6^ live cells. Samples were resuspended in PBS plus PTI and stained with Ghost Dye Red 780 (Tonbo Biosciences) for 30 minutes at 4°C, protected from light. After being washed with flow buffer plus PTI, samples were incubated with anti-CD16/32 (BioLegend or Tonbo Biosciences) for 5 minutes at room temperature to reduce nonspecific binding. Samples were stained with surface antibodies in brilliant stain buffer (BD Biosciences) for 30 minutes at 4°C, protected from light. For samples including intracellular markers, cells were washed with flow buffer plus PTI and fixed with Fixation/Permeabilization buffer (eBioscience) for 30 minutes at 4°C, protected from light. After being washed with 1X permeabilization buffer, samples were stained with intracellular antibodies in permeabilization buffer overnight at 4°C, protected from light. The following day, samples were washed once with permeabilization buffer and once with flow buffer. Samples were acquired on an Attune NxT flow cytometer (Thermo Fisher). The following surface antibodies were used: CD45 FITC or BV510 or BV605, CD3 PE-Cy5, NK1.1 BV510 or PE-Dazzle 594 or BV605, CD8α BV605 or APC-R700 or BV711, CD4 AlexaFluor700 or PE or FITC, CD19 PE-Dazzle 594 or APC, CD11b BD Horizon V450 or BD Horizon BB700, CD11c AlexaFluor700, F4/80 PE-Cy7, Ly-6G PE, CD107a BV421, PD-1 BV421, LAG-3 BV711, TIM-3 APC, TIGIT BV605, CD69 BV510 (BioLegend or BD Biosciences). The following intracellular antibodies were used: Perforin PE, IFNγ PE-Cy7 or BV711, GzmB APC (all from BioLegend). Two to four experimental repeats were completed to characterize immune infiltrate populations, activation, effector function, and exhaustion for both B78 and MC38 tumors across time via flow cytometry.

### Flow cytometry data analysis

Data were standardized between timepoints using rainbow beads (Spherotech). Data were analyzed using FlowJo (v10.7.1) and gates were determined using fluorescent minus one controls. At least one C57BL/6 non-mCherry mouse was included in each study to assist in setting mCherry gates.

### scRNAseq sample preparation and acquisition

Details on sample preparation and acquisition can be found at Erbe *et al.*^19^

### scRNAseq analysis of CD8 T, CD4 T, NK, and B78 tumor cells from B78 tumors treated with radio-immunotherapy

We utilized our scRNAseq data of B78 tumors, generated from an independent study,^19^ to evaluate metabolic genes and pathways in CD8 T, CD4 T, NK, and B78 tumor cells from vehicle and treated tumors. We used *Sox10* to select out the tumor cells, *Cd4* for CD4+ T cells, *Cd8α* for CD8+ cells, and *Ncr1* for NK cells for further clustering to determine subsets within each compartment. To account for potential technical variance across samples, we used a fuzzy clustering-based integration method (Harmony method).^106^ Downstream analysis for CD8 T cells and B78 tumor cells were based on Seurat single-cell analysis package^107^ including: graph-based clustering based on 20 Principal Components using *FindCluster* with different resolution from 0.1 to 2 to determine the number of clusters based on representative markers overlaid in the hierarchical tree across different resolution (Clustree R package), differential expression analysis using MAST^108^ implemented in Seurat with the cutoff average log2FC 0.25, adjusted *p*-value cutoff of 0.01, and at least 20% of cells expressing the markers. Visualization with UMAPs, heatmaps, dot plots, and violin plots was done using Seurat in R (v4.4.2) and the Bioconductor platform. Metabolism pathway and immune functional signature was calculated as the average of the genes in the Mouse Kyoto Encyclopedia of Genes and Genomes (KEGG)^75–77^ metabolism for each cell subset.

### Multiplex immunofluorescence sample preparation and acquisition

All multiplex immunofluorescence experiments were completed in CD8-mCherry mice with C57BL/6J tissues included as controls. To characterize CD8-mCherry mice, untreated B78 tumors and spleens were excised, formalin fixed, and paraffin-embedded (FFPE) for antibody staining with a panel of fluorescent markers (CD4, CD8α, mCherry) (all from Abcam) (**Figure S1**). Embedded sections were deparaffinized and hydrated prior to 7 min microwave antigen retrieval using 10X AR9 (Akoya Biosciences) diluted 1:10 with dH_2_O, a high pH antigen retrieval buffer, and placement in 1X Antibody Diluent / Block Solution (Akoya Biosciences) for 10 min at RT. Next, primary antibodies were sequentially applied upon removal of blocking solution at the following dilutions and incubation times: CD4 – 1:500 for 15 min, CD8α –1:250 for 15 min, mCherry 1:500 – for 10 min. Following 1X tris-buffered saline washes, ∼200 μL diluted rat or rabbit secondary antibody (Abcam) was applied to each slide and incubated for 10 min. Following a 1X tris-buffered saline wash, Opal dyes were applied at 1:100 dilution for 10 min in 1X Amplification Diluent (Akoya Biosciences): (tumor and spleen sections) CD4 – Opal-dye 520, CD8α – Opal dye 620, mCherry – Opal dye 570 (all from Akoya Biosciences). Finally, stained sections were incubated in diluted DAPI (Akoya Biosciences) for 10 min at RT for nuclear labeling and mounted on glass coverslips with Prolong Gold Antifade Reagent (Invitrogen) for imaging. Imaging was performed on whole tumor and spleen slides at 4× and then regions of interest (ROIs) were selected and imaged at 20× using a Vectra multispectral imaging system (Akoya Biosciences). One experimental replicate was performed.

For all other studies using treated and vehicle CD8-mCherry mice, excised B78 tumors were formalin fixed and paraffin-embedded for antibody staining with a panel of fluorescent markers (perforin, SOX10, CD161, cPARP-1, CD4, CD8α) (all from Abcam). Two tumors were embedded/paraffin block. Excised spleens and lymph nodes from B78 tumor bearing mice were formalin fixed and processed into FFPE blocks. 2.5 mm cores were punched from these donor blocks to create a tissue microarray (TMA) to fit samples from this study cohort. This mouse TMA was stained with a panel of fluorescent markers (F4/80, CD69, CD19, CD3, CD4, CD8α) (all from Abcam) (**Figures 7, S6**). Embedded sections were deparaffinized and hydrated prior to 7 min microwave antigen retrieval using 10X AR9 (Akoya Biosciences) diluted 1:10 with dH_2_O, a high pH antigen retrieval buffer, and placement in 1X Antibody Diluent / Block Solution (Akoya Biosciences) for 10 min at RT. Next, primary antibodies were sequentially applied upon removal of blocking solution at the following dilutions and incubation times at RT: (tumor sections) perforin – 1:100 for 30 min, SOX10 – 1:100 for 60 min, CD161 – 1:4000 for 5 min, cPARP-1 – 1:100 for 15 min, CD4 – 1:500 for 15 min, CD8α –1:250 for 15 min; (TMA sections) F4/80 – 1:50 for 45 min, CD69 – 1:50 for 30 min, CD19 – 1:500 for 15 min, CD3 – 1:500 for 15 min, CD4 – 1:500 for 15 min, CD8α – 1:250 for 15 min. Following 1X tris-buffered saline washes, ∼200 μL diluted rat or rabbit secondary antibody (Abcam) was applied to each slide and incubated for 10 min. Following a 1X tris-buffered saline wash, Opal dyes were applied at 1:100 dilution for 10 min in 1X Amplification Diluent (Akoya Biosciences): (tumor sections) perforin – Opal dye 520, SOX10 – Opal dye 540, CD161 – Opal dye 570, cPARP-1 – Opal dye 620, CD4 – Opal-dye 650, CD8α – Opal dye 690; (TMA sections) F4/80 – Opal dye 520, CD69 – Opal dye 540, CD19 – Opal dye 570, CD3 – Opal dye 620, CD4 – Opal dye 650, CD8α – Opal dye 690 (all from Akoya Biosciences). Finally, stained sections were incubated in diluted DAPI (Akoya Biosciences) for 10 min at RT for nuclear labeling and mounted on glass coverslips with Prolong Gold Antifade Reagent (Invitrogen) for imaging. Imaging was performed on whole tumor slides at 4× and then regions of interest (ROIs) were selected and imaged at 20× using a Vectra multispectral imaging system (Akoya Biosciences) – 8 ROIs/tumor, 5 at tumor exterior and 3 at tumor interior. Imaging was performed on the TMA by taking a 3×3 grid within each spleen or lymph node core, at 20× using a Vectra multispectral imaging system (Akoya Biosciences). A spectral library was also generated to separate spectral curves for each of the fluorophores. One experimental replicate was performed.

### Multiplex immunofluorescence data analysis

Resulting multispectral images were analyzed using Nuance and inForm software (Akoya Biosciences). A spectral library algorithm was created using multispectral image cubes from Vectra to define distinctive spectral curves for each fluorophore, chromogen, and counterstain to adjust for background effects and accurately quantify positive staining of biomarkers using inForm software. InForm software analysis allows objective counting of cell populations and biomarkers and increases the accuracy of the statistical analysis. Algorithms for subcellular compartment separation were created by machine learning to identify nuclei and membrane, respectively, to accurately assign associations for positive staining to a specific compartment in the tumor microenvironment. To create each algorithm, 10% of the total image dataset was used for each experiment. Positivity thresholds were set for each antibody in each tissue type (tumor, spleen, lymph node).

### Cryo-fluorescence tomography imaging

B78-bearing CD4 mCherry^59^ mice were treated with RT, Hu14.18-IL2, and αCTLA-4 or no treatment. On Day 12 post-treatment mice were euthanized. Hexanes were added to a tall dewer filled with dry ice. Euthanized mice were fully submerged in hexanes to slowly cool mice for ∼15 min and then were stored at -80°C. Mice were then frozen onto a single block of O.C.T. (Ultrafreeze OCT Clear, Cancer Diagnostics Durham, NC, USA) that was then sectioned into 20 µm slices over 17 h at −15°C (Xerra™, Emit Imaging, Baltimore, MD). Imaging was performed on the Emit Imaging Xerra platform, an automated cryo-fluorescence tomography system with an integrated cryo-microtome and proprietary software that acquires 2D white light and fluorescence images of serial sections and reconstructs them into 3D volumes. For each section, brightfield images were acquired sequentially along with images acquired using at 555 nm excitation wavelength and a 586/15 nm detection filter of for mCherry. Images were analyzed using ImageJ.

### Statistical analysis

All details on statistical tests, replicates, etc. can be found within the figure captions.

Mice survival probabilities were estimated using the Kaplan-Meier method. Pairwise log-rank tests with Benjamini-Hochberg adjustment were used to compare survival curves of the different treatment groups within each experiment. Tumor volumes were considered "missing" after mouse death. All remaining available data was used for the tumor-growth analysis. Linear mixed models accounting for mouse id as a random intercept were fit for each experiment, with treatment, time in days, and the interaction of treatment & day included as fixed effects. The outcome, tumor volume, was transformed by log_10_(y+0.0001) to satisfy model assumptions. The small constant (0.0001) was added to handle tumor volumes of 0. The tumor-growth rates of the treatment groups within each experiment were compared using pairwise contrasts and adjusted for multiple comparisons using Tukey’s method. Differences in primary and rechallenge cures were calculated via Fisher exact test. Mouse analyses were carried out in R v4.1.2.

Flow cytometry results are represented as scatter dot plots showing mean (bar) ± standard deviation. Each dot represents a single mouse tumor. A one-way ANOVA was calculated with Šίdák’s multiple comparisons test, α = 0.05 (GraphPad Prism v10.4).

*In vivo* metabolic imaging results are represented as dot plots showing mean ± standard deviation, stratified by treatment group and imaging day. Each dot represents a single tumor or CD8 T cell. A multi-level linear mixed effects model with animal specific random effects was used to evaluate differences in metabolism parameters on single-cell levels between treatment groups and imaging day assessment time points. Treatment group, day, and the interaction effect between day and treatment groups were included as predictor variables. A compound symmetry correlation structure was used to account for correlations between subsamples within each animal. Model assumptions were verified by examining residual plots. All reported *p*-values are two-sided and *p*<0.05 was used to define statistical significance. Statistical analyses were conducted using SAS software v9.4 (SAS Institute, Cary NC). Effect size was also calculated to assess differences in optical metabolic imaging parameters between treatment groups using Glass’s Δ because comparisons of very large sample sizes of individual cells nearly always pass traditional significance tests unless the population effect size is truly zero. Glass’s Δ is defined as: (mean experimental group – mean control group) ∕ standard deviation control group. A Glass’s Δ > 0.8 was chosen to indicate significant effect size based on previous studies.^59,122,123^

*In vitro* metabolic imaging results are represented as dot plots showing mean ± standard deviation. Each dot represents a single tumor cell. Mann–Whitney statistical tests for non-parametric, unpaired comparisons were performed to assess differences in metabolic parameters between treatment groups (GraphPad Prism v10.4). Effect size was also calculated.

Single-cell RNA sequencing results are represented as UMAPs, heatmaps (relative gene expression), dot plots (average gene expression), and violin plots showing mean ± standard deviation. Differences between specific metabolic genes across cell clusters were calculated via Wilcoxon rank-sum test with Benjamini-Hochberg correction (R v4.4.2).

Multiplex immunofluorescence results are represented as scatter dot plots showing mean (bar) ± standard deviation, where each dot represents a single field of view (FOV). Data is also represented as line plots showing the average value across several FOV. A one-way ANOVA was calculated with Šίdák’s multiple comparisons test, α = 0.05 (GraphPad Prism v10.4).

## Data availability

- scRNAseq data have been deposited at Gene Expression Omnibus under accession number: GSE310495 reported within Erbe *et al.*^19^
- Original code used to analyze scRNAseq data can be found at the following GitHub repository: https://github.com/huydinhlab reported within Erbe *et al.*^19^
- The raw OMI data supporting this article can be found at the following GitHub repository: https://github.com/skalalab/heaton_a-B78-MC38-invivo-data

## Acknowledgements

This work was supported by a Morgridge Interdisciplinary Postdoctoral Fellowship to A.R.H; The Midwest Athletes against Childhood Cancer Fund; Stand Up to Cancer, The St. Baldrick’s Foundation; The Crawdaddy Foundation; The Cancer Research Institute; The Alex’s Lemonade Stand Foundation; The End Kids Cancer Foundation; The Hartwell Foundation; The Carol Skornicka Chair in Biomedical Imaging; Retina Research Foundation Daniel M. Albert Chair; and the National Institutes of Health [R35CA197078, 5P30CA014520-40, UL1TR002373, P01CA250972, U54CA232568, R01CA278051, R01CA272855]. The authors acknowledge the UW Small Animal Imaging & Radiotherapy Facility and the UWCCC Flow Cytometry Laboratory, both supported by P30 CA014520. The authors acknowledge the UW Translational Research Initiatives in Pathology laboratory, supported by the UW Department of Pathology and Laboratory Medicine, UWCCC (P30 CA014520) and the Office of The Director – NIH (S10 OD023526) for use of their facilities and creation of a TMA (using Grand Master Microarrayer S10 OD023526). The authors thank Mohammed Farhoud and Emit Imaging (cryo-fluorescence tomography imaging), Sritha Moram (mouse assistance), Wenxuan Zhao (data analysis), Jens Eickhoff and Meredith Hyun (statistical analysis), Matt Stefely (figure formatting), and Alicia Williams (editing).

## Corresponding author

Requests for resources and reagents should be directed to Melissa Skala.

## Author contributions

Conceptualization, A.R.H., P.M.S., M.C.S., A.K.E., A.L.R.; Methodology, A.R.H., A.G., A.H., A.S.F., A.K.E.; Investigation, A.R.H., N.J.B., A.G., A.H., A.S.F., S.K.B., M.A.F, N.W.T., D.V.S., G.M.L., A.A.H., A.D., A.K.E.; writing—original draft, A.R.H.; writing—initial review & editing, P.M.S. and M.C.S.; final review and editing-all authors; funding acquisition, A.R.H., P.M.S., M.C.S.; resources, H.Q.D.; supervision, A.R.H., H.Q.D., P.M.S., M.C.S.

