## Supplemental Figures and Table for "Divergent role of CD8 T cells with distinct metabolic phenotypes during curative radio-immunotherapy in hot versus cold tumors"

**Supplemental Information**

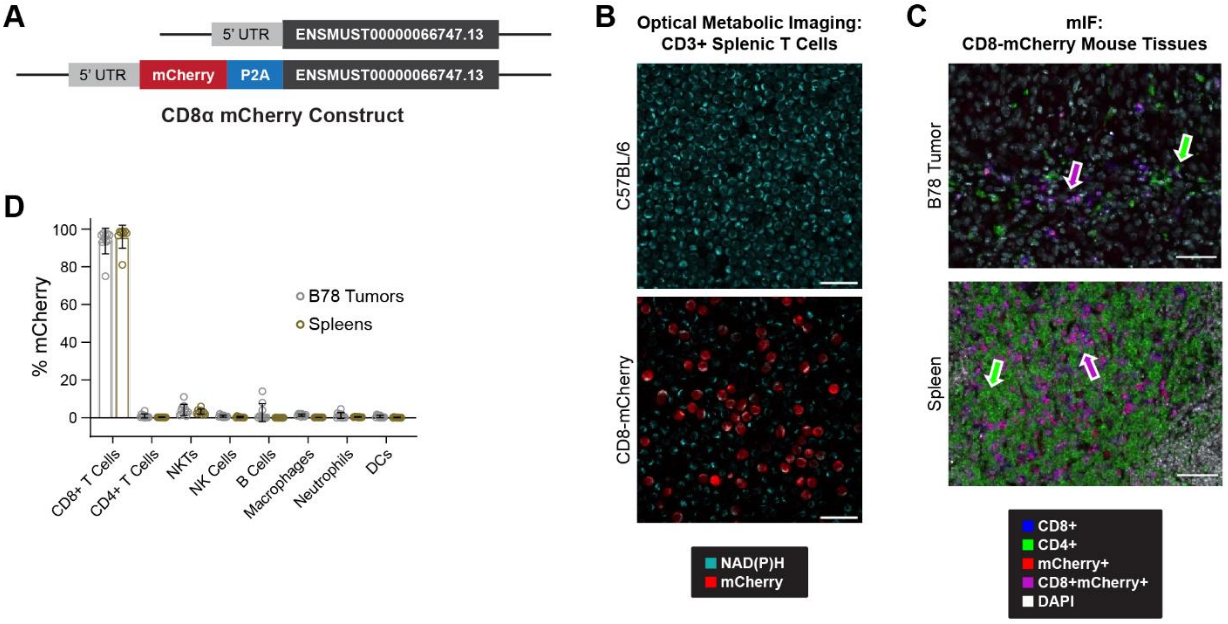


**Figure S1. Characterization of CD8-mCherry reporter mouse shows high specificity of mCherry label to CD8-T cells**

**A)** Construct used to knock-in mCherry near CD8α allele via CRISPR/Cas 9 to create CD8-mCherry reporter mouse. **B)** CD3+ splenic T cells isolated from wild type C57BL/6J mouse (top) and CD8-mCherry mouse (bottom) and imaged via OMI. All T-cells contain NAD(P)H (cyan) while only the reporter mouse expresses mCherry in ~50% of the cells (red) (Scale bar 25 µm). **C)** Multiplex immunofluorescence images of untreated reporter mouse B78 melanoma tumor and spleen show that mCherry is only expressed on cells that co-express CD8 (purple). There is no free mCherry signal (red). There is free CD4 signal (green) as this cell type does not express mCherry (Scale bar 50 µm). **D)** Flow cytometry of untreated B78 melanoma tumors and spleens shows that nearly 100% of CD8-T cells express mCherry while all other immune cells express minimal to no mCherry (*n* = 10 B78 tumors, *n* = 8 spleens, mean ± SD, 2 experimental repeats). Characterization across all techniques confirmed the mCherry knock-in was highly specific to CD8α allele providing confidence in subsequent experiments investigating mCherry+ populations.

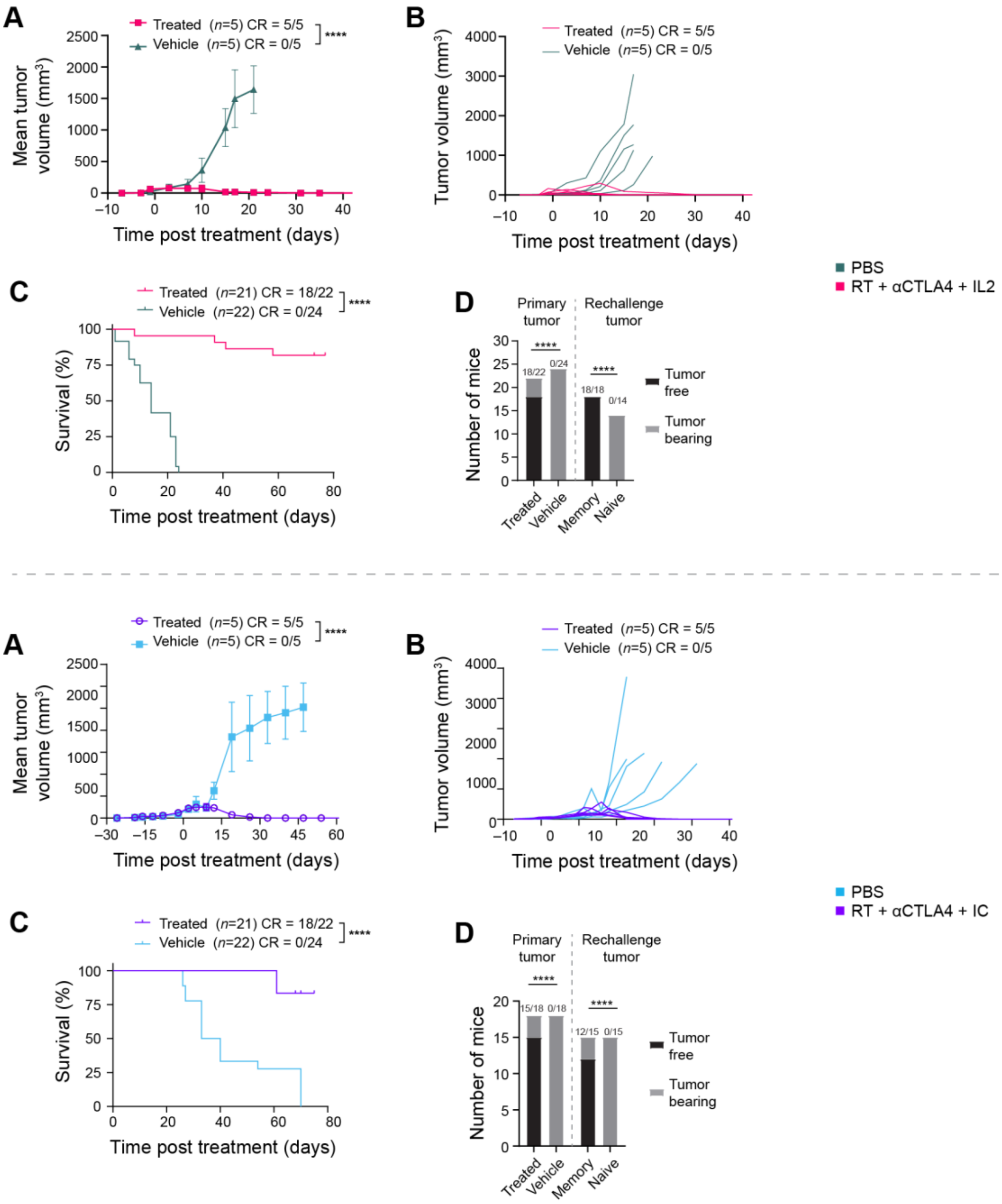


**Figure S2. MC38 and B78 response to radio-immunotherapy repeated in C57BL/6J wild type mice**
**A)** Mean tumor-growth curves for treated and vehicle C57BL/6J mice bearing MC38 tumors (pairwise contrasts, *n* = 5 mice/group, mean ± SEM, 1 representative experiment shown, 3 experimental repeats). **B)** Individual tumor-growth curves for treated and vehicle C57BL/6J mice bearing MC38 tumors (*n* = 5 mice/group, 1 representative experiment shown, 3 experimental repeats). **C)** Kaplan-Meier survival curve showing % survival of treated and vehicle C57BL/6J mice bearing MC38 tumors (pairwise log-rank test, *n* = 46 mice, 3 experimental repeats combined). **D)** Number of tumor free C57BL/6J mice from treated and vehicle primary MC38 tumors (Fisher exact test, *n* = 46, 3 experimental repeats) and mice from treated and naïve MC38 tumor rechallenge 2 months later (Fisher exact test, *n* = 32, 3 experimental repeats). **E)** Mean tumor-growth curves for treated and vehicle C57BL/6J mice bearing B78 tumors (pairwise contrasts, *n* = 5 mice/group, mean ± SEM, 1 representative experiment shown, 3 experimental repeats). **F)** Individual tumor-growth curves for treated and vehicle C57BL/6J mice bearing B78 tumors (*n* = 5 mice/group, 1

representative experiment shown, 3 experimental repeats). **G)** Kaplan-Meier survival curve showing % survival of treated and vehicle C57BL/6J mice bearing B78 tumors (pairwise log-rank test, $n = 36$ mice, 3 experimental repeats combined). **H)** Number of tumor free C57BL/6J mice from treated and vehicle primary B78 tumors (Fisher exact test, $n = 36$, 3 experimental repeats) and mice from treated and naïve B78 tumor rechallenge 2 months later (Fisher exact test, $n = 30$, 3 experimental repeats) (*$p < 0.05$, **$p < 0.01$, ***$p < 0.001$, ****$p < 0.0001$).

**A**

| Population | Gating Phenotype (after Cells/Single/Live) |
|---|---|
| CD45 Lymphocytes | CD45+ |
| CD8 T Cells | CD45+/CD3ε+/NK1.1-/CD8α+ |
| CD4 T Cells | CD45+/CD3ε+/NK1.1-/CD4+ |
| NKT Cells | CD45+/CD3ε+/NK1.1+ |
| NK Cells | CD45+/CD3ε-/NK1.1+ |
| B Cells | CD45+/CD3ε-/NK1.1-/CD19+ |
| Macrophages | CD45+/CD3ε-/Ly6G-/F4/80+ |
| Neutrophils | CD45+/CD3ε-/Ly6G+/CD11b+ |
| DCs | CD45+/CD3ε-/Ly6G+/CD11b+/CD11c+ |

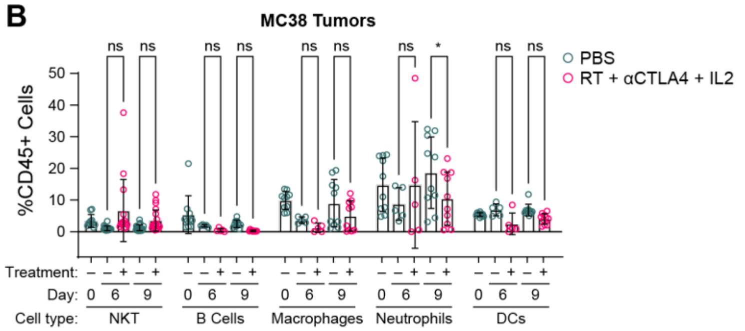


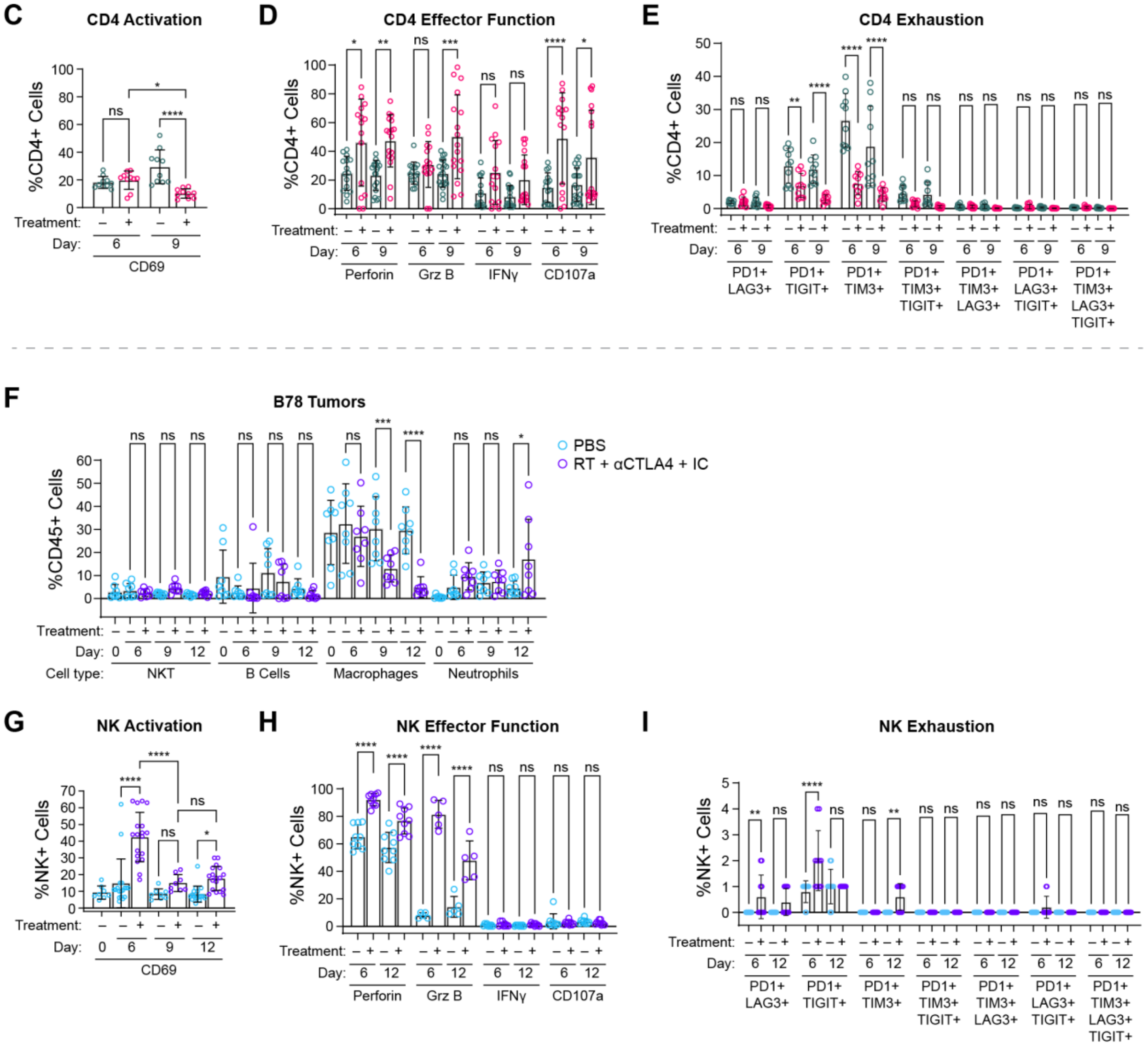


**Figure S3. Flow cytometry gating strategies with additional MC38 and B78 flow data**

**A)** Flow cytometry gating phenotype used for all immune cell populations in Figures 1,3, and S3. **B)** Flow cytometry of NKT, B, macrophages, neutrophils, and DCs from treated and vehicle MC38 tumor bearing mice across treatment time (one-way ANOVA, *n* = 5-20 mice/group, 2-4 experimental repeats, mean ± SD, % CD45+ cells). **C-E)** Flow cytometry of CD4 activation, effector function, and exhaustion from treated and

vehicle MC38 tumor bearing mice across treatment time. **F)** Flow cytometry NKT, B, macrophages, and neutrophils from treated and vehicle B78 tumor bearing mice across treatment time (one-way ANOVA, $n$ = 10 mice/group, 2 experimental repeats, mean ± SD, % CD45+ cells). **G-I)** Flow cytometry of NK cell activation, effector function, and exhaustion from treated and vehicle B78 tumor bearing mice across treatment time (*$p < 0.05$, **$p < 0.01$, ***$p < 0.001$, ****$p < 0.0001$).
**C-E)** $n$ = 10-20 mice/group, one-way ANOVA, mean ± SD, % CD4+ cells, 2-4 experimental repeats.
**G-I)** $n$ = 10-20 mice/group, one-way ANOVA, mean ± SD, % NK+ cells, 2-4 experimental repeats.

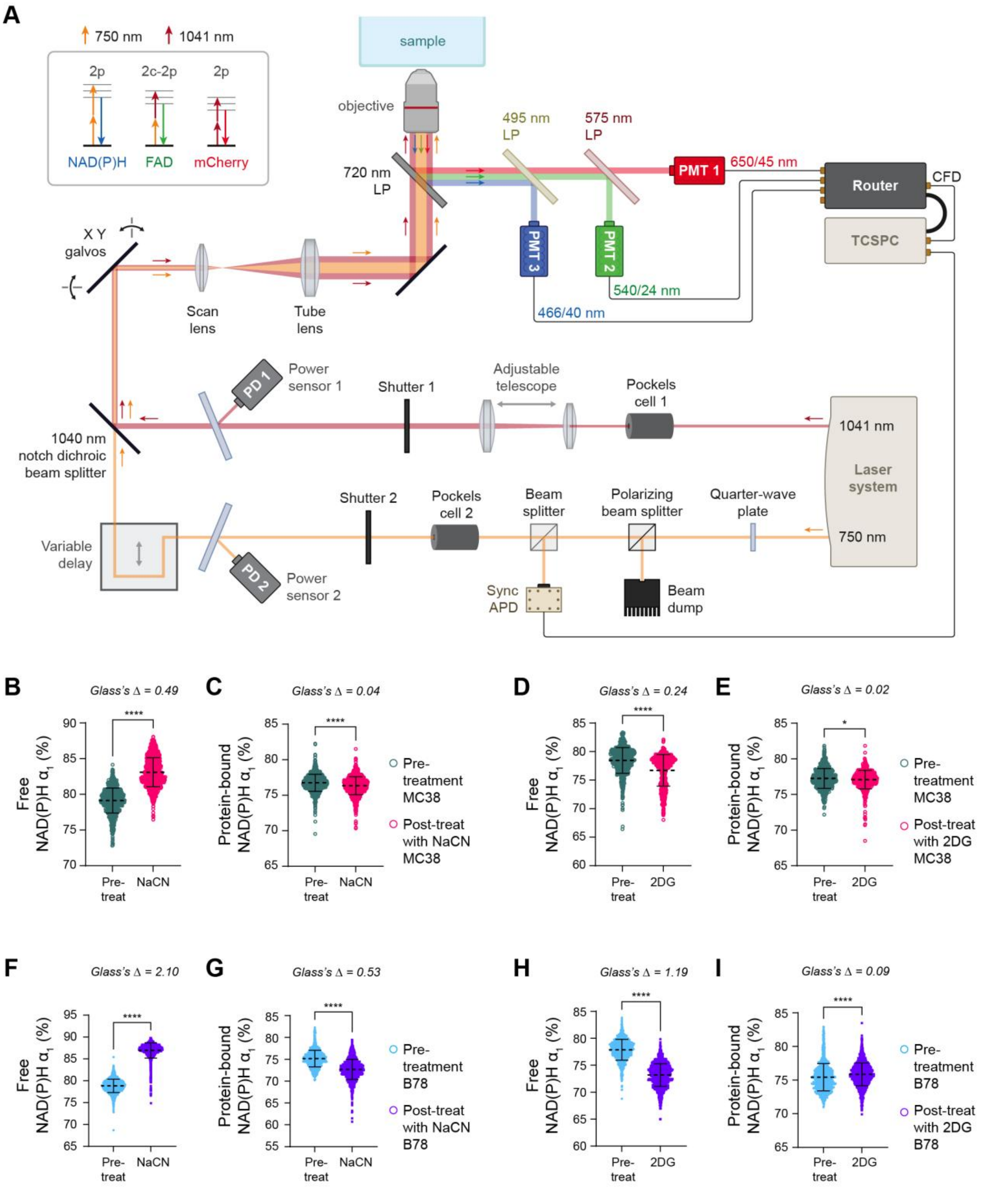


**Figure S4. Optical metabolic imaging system and *in vitro* MC38 and B78 metabolic changes with electron transport chain (NaCN) and glycolysis (2DG) metabolic inhibitor treatments. A)** Two-photon imaging system scheme including 750 and 1041 nm lasers, three photo-multiplier tube detectors, and single photon counting capabilities. Jablonski diagram shows how wavelength mixing with 750 nm laser and 1041 nm laser excites NAD(P)H, FAD, and mCherry simultaneously. **B-C)** Quantified single cell MC38 metabolic changes with NaCN treatment ($n$ = 932 pre-treatment cells, $n$ = 1154 post-treatment NaCN cells). **See Table S1** for parameter definitions. **D-E)** Quantified single cell MC38 metabolic changes with 2DG treatment ($n$ = 643 pre-treatment cells, $n$ = 612 post-treatment 2DG cells). **F-G)** Quantified single cell B78 metabolic changes with NaCN treatment ($n$ = 1488 pre-treatment cells, $n$ = 1393 post-treatment NaCN cells). **H-I)** Quantified single cell B78 metabolic changes with 2DG treatment ($n$ = 1113 pre-treatment cells, $n$ = 1589 post-treatment 2DG cells)

**B-I)** 2 technical replicates, 2 experimental repeats, unpaired T test *$p < 0.05$, ****$p < 0.0001$, Glass's Δ, mean ±SD.

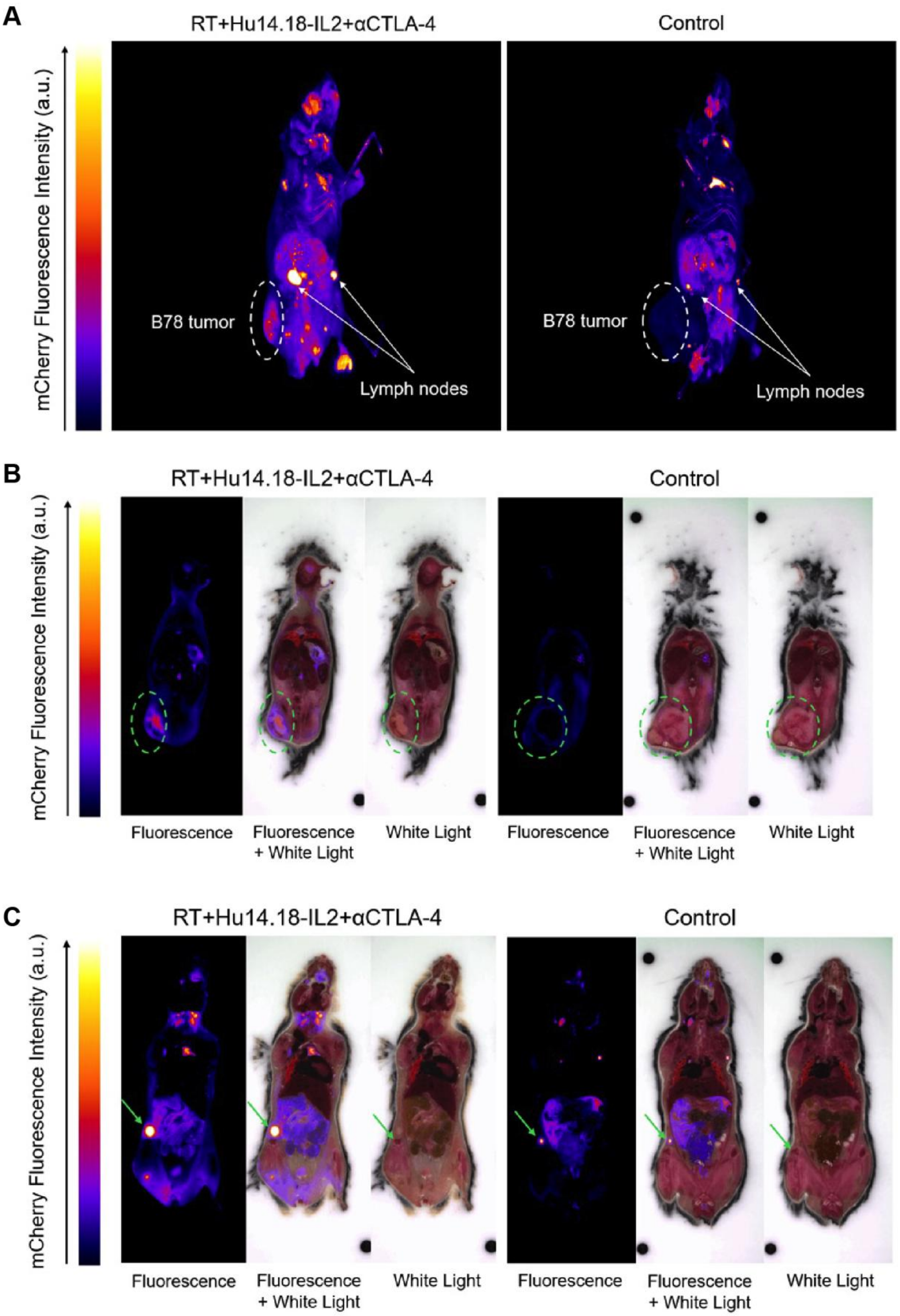

A
RT+Hu14.18-IL2+αCTLA-4
Control
mCherry Fluorescence Intensity (a.u.)
B78 tumor
Lymph nodes
B78 tumor
Lymph nodes
B
RT+Hu14.18-IL2+αCTLA-4
Control
mCherry Fluorescence Intensity (a.u.)
Fluorescence
Fluorescence + White Light
White Light
Fluorescence
Fluorescence + White Light
White Light
C
RT+Hu14.18-IL2+αCTLA-4
Control
mCherry Fluorescence Intensity (a.u.)
Fluorescence
Fluorescence + White Light
White Light
Fluorescence
Fluorescence + White Light
White Light

**Figure S5. Cryo-fluorescence tomography imaging of B78-bearing mice illustrates increased tumor and lymph node infiltration of mCherry+ T cells after radio-immunotherapy compared to vehicle control. A)** 3D volumetric reconstruction of mCherry T cell mice[66] bearing B78 melanoma tumors. Mice treated with RT, Hu14.18-IL2, and αCTLA-4 had increased T cell infiltration into the tumor (white dashed circle) and inguinal lymph nodes (white arrows) compared to vehicle control mice. Relative mCherry intensity from mCherry expressing CD8, CD4, NKT cells[66]. **B-C)** Coronal sections of mCherry T cell bearing B78 melanoma tumors via fluorescence images alone, fluorescence overlaid on white light images, or white light images alone. The coronal sections in **B** intentionally include the greatest cross section of tumor (green dashed circle), while the coronal sections in **C** intentionally include the greatest cross section of tumor draining lymph node (green arrow). Mice treated with radio-immunotherapy show the presence of mCherry+ T cell signal within the tumor (green dashed circle) while control mice do not. Both treated and control mice show the presence of mCherry+ T cell signal within the tumor draining lymph node (green arrow), though the treated mouse expresses more signal intensity and area.

**A-C)** One experimental repeat performed with Emit Imaging. Full videos can be found here: https://github.com/skalalab/heaton_a-B78-MC38-invivo-OMI-data

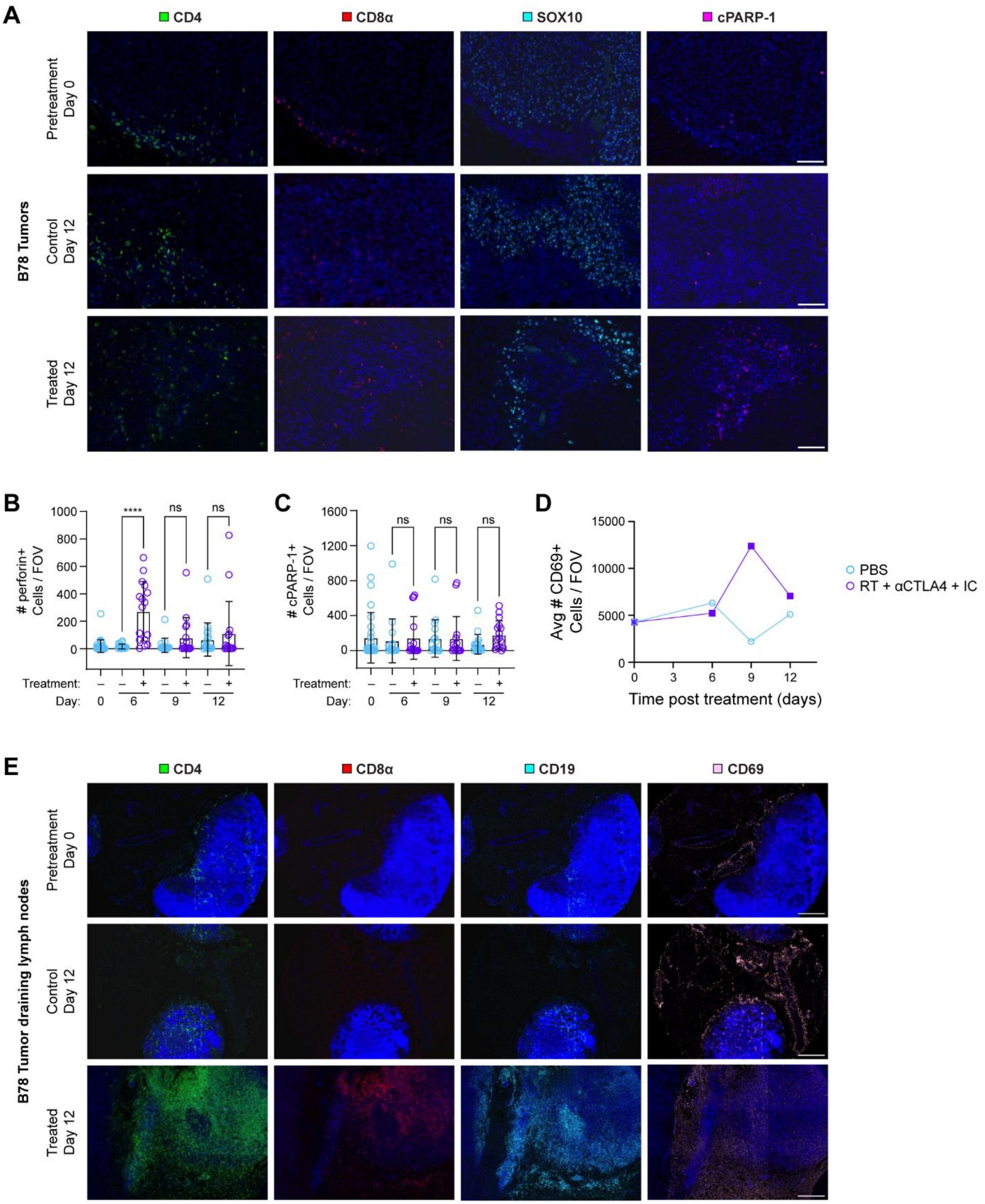


**Figure S6. Single channel B78 tumor and lymph node immunofluorescence images with additional quantified data**

**A)** 20× single channel B78 tumor immunofluorescence images at day 0 and 12 of treatment (*n* = 2-4 mice/group, 1 experimental replicate, scale bar 100 μm). **B-C)** Quantified changes within B78 tumor images of perforin and cPARP-1 (*n* = 2-4 mice/group, *n* = 8 images/tumor, 1 experimental replicate, one-way ANOVA, mean ± SD). **D)** Quantified CD69+ cells from lymph nodes from treated and vehicle B78 tumor bearing mice at day 0 and 12 of treatment (*n* = 1-2 mice/group, *n* = 2-4 lymph node/group, 1 experimental replicate, average shown). **E)** Single channel immunofluorescence images from B78 tumor-draining lymph nodes at day 0 and 12 of treatment. 3×3 20× images stitched together/lymph node (*n* = 1-2 mice/group, *n* = 9 images/lymph node, 1 experimental replicate, scale bar 300 μm) (* $p < 0.05$, ** $p < 0.01$, *** $p < 0.001$, **** $p < 0.0001$).

**Table S1. Fluorescence lifetime imaging and morphology parameters.**

| **Parameter** | **Description** | **Included to classify CD8 T cells** | **Included to classify tumor cells** |
|---|---|---|---|
| NAD(P)H $\tau_1$ | Free NAD(P)H lifetime | ✓ | ✓ |
| NAD(P)H $\tau_2$ | Protein-bound NAD(P)H lifetime | ✓ | ✓ |
| NAD(P)H $\alpha_1$ | Proportion of free NAD(P)H | ✓ | ✓ |
| NAD(P)H $\alpha_2$ | Proportion of protein-bound NAD(P)H | -- | -- |
| NAD(P)H $\tau_m$ | NAD(P)H mean lifetime: $\alpha_1 \tau_1 + \alpha_2 \tau_2$ | ✓ | ✓ |
| FAD $\tau_1$ | Protein-bound FAD lifetime | ✓ | ✓ |
| FAD $\tau_2$ | Free FAD lifetime | ✓ | ✓ |
| FAD $\alpha_1$ | Proportion of protein-bound FAD | ✓ | ✓ |
| FAD $\alpha_2$ | Proportion of free FAD | -- | -- |
| FAD $\tau_m$ | FAD mean lifetime: $\alpha_1 \tau_1 + \alpha_2 \tau_2$ | ✓ | ✓ |
| Fluorescence lifetime redox ratio | FLIRR: NAD(P)H $\alpha_2$ / FAD $\alpha_1$ | ✓ | ✓ |
| Area | Cell area | ✓ | -- |
